# Genomic and morphometric evidence for Austronesian-mediated pig translocation in the Pacific

**DOI:** 10.1101/2025.07.07.663491

**Authors:** David W. G. Stanton, Aurelie Mannin, Allowen Evin, Kristina Tabbada, Anna Linderholm, Rosie Drinkwater, Olaf Thalmann, Said I Ng’Ang’A, Noel Amano, Atholl Anderson, Ross Barnett, Patrick Barrière, Stuart Bedford, Peter Bellwood, Adam Brumm, Trung Cao Tien, Geoffrey Clark, Richard Crooijmans, Thomas Cucchi, Michelle S. Eusebio, Linus G Flink, Peter Galbusera, Martien Groenen, Budianto Hakim, Stuart Hawkins, Holly Heiniger, Kristofer M Helgen, Michael J. Herrera, Terry Hunt, Andrew C Kitchener, Carol Lee, Alastair A Macdonald, Hendrik-Jan Megens, Erik Meijaard, Kieren J. Mitchell, Christopher Moran, Karen Mudar, Karma Nidup, Marc Oxenham, Rinzin Pem, Philip Piper, Kyle Schachtschneider, Lawrence Schook, Pradeepa Silva, Matthew Spriggs, Samuel Turvey, Una Strand Viðarsdóttir, Murray P. Cox, Tim Denham, Jamie Gongorra, Keith Dobney, Greger Larson, Laurent A. F. Frantz

## Abstract

Human mediated translocation of non-native pig species (genus *Sus*) to the islands of Wallacea and Oceania has significantly altered local ecosystems. To investigate the timing and trajectory of these introductions, we conducted both genomic analyses of 576 pig nuclear genomes and a geometric morphometric analysis of 714 modern and ancient dental remains. Our analyses demonstrate that feral and domestic pigs in Wallacea and Oceania possess diverse ancestries resulting from the introduction of multiple, sequential pig populations followed by gene flow. Despite the variability in their genomic ancestry these pigs all possess a distinct tooth morphology, and a genetic link to the Chinese domestic pig populations that accompanied the dispersal of Austronesian language speakers ∼4,000-3,000 years ago via Taiwan and the Philippines.

## Main Text

People have both involuntarily (e.g., commensal species such as rats and mice) and deliberately (e.g., as game management or livestock) transported animal species beyond their native ranges for millennia. These introductions have often substantially altered local ecosystems, particularly on islands (*1*). The translocation of vertebrates eastwards across the Wallace Line (Fig. 1), into Wallacea, a biogeographic region between the Asian and Australasian biotas, has had dramatic environmental impacts (*2*). Notably, while the natural range of the genus *Sus* (pigs) is primarily west of the Wallace Line, and limited to the westernmost part of Wallacea (i.e. Sulawesi and nearby islands), pigs are now widespread across archipelagoes in Wallacea, Melanesia, Micronesia, and Polynesia (Fig. 1A).

**Fig. 1.**
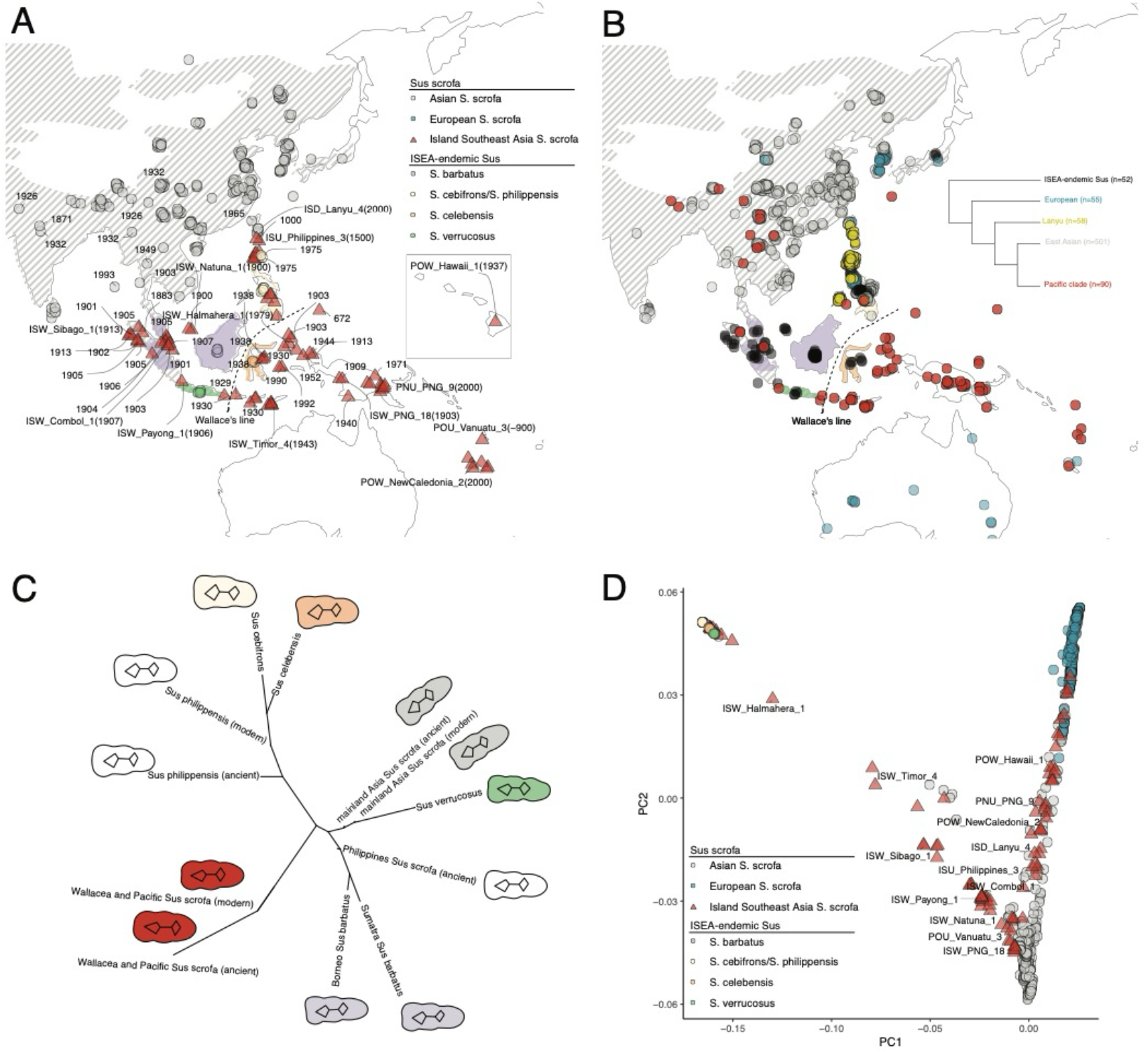
(**A) Map of sampling locations**. Geographical distribution of ancient and modern S. scrofa samples, with dates, used in this study. **(B) Mitochondrial analysis**. Geographic distribution of major mtDNA haplogroups based on data from ∼705 control regions and ∼323 whole mitochondrial genome data (see Figure S1 for full mitogenome tree). **(C) Geometric morphometrics**. Neighbour-joining network displaying morphometric similarities in the third lower molar shape between S. scrofa and ISEA-endemic Sus species. Tooth outlines represent population mean shapes. **(D) Principal component analysis (PCA)**. PCA plot illustrating the genetic variation of Eurasian S. scrofa and related species.

The human-mediated translocation of pigs east of the Wallace Line most likely began in pre-Neolithic times, >4,000 years ago, prior to the dispersal of domestic animals into the region (*3*). The recent discovery on Sulawesi of the world’s oldest figurative artwork (>50,000 years old) that depicts a local endemic warty pig species, *Sus celebensis*, demonstrates the deep-rooted relationship between humans and pigs in the region (*4, 5*). Mitochondrial DNA and zooarcheological analyses have identified *S. celebensis* remains from pre-Neolithic layers (older than 4,000 years) in Liang Bua cave, Flores (*6*–*8*), although these have not yet been directly radiocarbon dated. The presence of this species in pre-Neolithic deposits, ∼300 km from its natural range in Sulawesi across the Flores Sea suggests these pigs were intentionally translocated by hunter-gatherers.

The first evidence for the presence of the Eurasian wild pig (*Sus scrofa*, the ancestor of domestic pigs), east of the Wallace line is also on Flores and occurred by ∼4,000 years ago (*6*–*8*).

Mitochondrial DNA analyses suggest that, as opposed to *S. celebensis*, the introduction of *S. scrofa* was a great deal more widespread and populations now exist as free-living populations in Wallacea, Melanesia, and Polynesia. (Fig. 1; (*6*)). These *S. scrofa* populations were likely first introduced 4,000-3,000 years ago during the expansion of Austronesian speakers from mainland East Asia and Taiwan, through the Philippines and Northern Wallacea, into Melanesia and Polynesia (*9*). This is supported by the remains of pigs (∼2,900 years before present [BP]) excavated in Melanesia (e.g., Vanuatu) from Lapita archaeological contexts associated with the Austronesian expansion (*10*–*12*).

Pigs from these regions predominantly display a particular tooth shape and a distinct mitochondrial haplogroup known as the “Pacific Clade” (*6, 13*). This mtDNA signature has also been identified in wild caught *S. scrofa* individuals from Java, Sumatra, and mainland Southeast Asia, including Laos, Vietnam, and Yunnan Province in southwestern China. This distribution suggests that the pigs introduced to Wallacea originated from peninsular Southeast Asia (*6, 13*). In contrast, modern domestic pigs found along the traditionally described route of the Austronesian expansion in Southeast China, Taiwan, and the Philippines, possess distinct Lanyu and East Asian mitochondrial signatures (*6, 13*). Combined with archaeobotanical evidence, these data suggest that Austronesian speakers only acquired pigs, and a variety of horticultural and arboreal crops (*14*–*17*), after leaving the Philippines, through interactions with local groups.

West of the Wallace Line, however, the “Pacific clade” mitochondrial haplogroup has only been identified in free-living pigs (*6, 13*). The absence of this haplogroup in modern domestic pigs thus challenges the idea that it represents a marker of domestic pigs introduced from mainland Asia. Furthermore, recent nuclear genome analyses have shown that, although *S. scrofa* populations are primarily found in mainland Eurasia, some *S. scrofa* populations naturally occur as wild boar in Western Indonesia (*18, 19*) (Sumatra, and Java; Fig. 1A), where the “Pacific Clade” haplogroup is also found (*6, 13*). These *S. scrofa* populations occur in sympatry with other, Island South East Asian (ISEA) endemic *Sus* species (ISEA-endemic *Sus*), found on Borneo, Sumatra (both *S. barbatus*), and Java (*S. verrucosus*). It thus remains unclear whether wild *S. scrofa* populations dispersed naturally, or were transported as wild individuals eastward across the Wallace Line.

The successful natural dispersal of the ancestor of another ISEA-endemic *Sus, S. celebensis*, from Borneo to Sulawesi (*18*), and its subsequent human-mediated translocation to the Lesser Sunda Islands suggests that a combination of natural and human-mediated dispersal of wild *S. scrofa* across the Wallace Line and onward toward Melanesia is plausible. In this scenario, it is conceivable that the pigs discovered in Lapita archaeological contexts were the descendants of domestic pigs introduced by Austronesian-speaking groups from mainland Asia, and that they acquired mitochondrial haplotypes belonging to the Pacific Clade through gene flow with other pig populations that were already present in Wallacea.

To address these questions and to establish the geographic origin of the pig populations in Wallacea, Melanesia, Micronesia and Polynesia, we sequenced 117 *Sus* nuclear genomes and generated geometric morphometric data (third lower molar) for 401 modern and 313 archaeological specimens. The genomic data generated here included 54 modern genomes, 60 historical genomes (1896-1993) and three ancient genomes including one each from the Philippines (∼1500 AD), Palau (Micronesia; ∼672 AD), and the Lapita culture site of Teouma on Efate Island, Vanuatu (∼900 BC). We analyzed these data alongside publicly available data that included 400 nuclear genomes (Fig. 1A) and ∼700 mtDNA control region sequences.

### The origin of the “Pacific Clade” mitochondrial haplogroup and associated morphometric signature

To assess the monophyletic strength of the Pacific Clade mitochondrial haplogroup using whole mitogenomes, we first built a maximum likelihood phylogeny using 585 mtDNA genomes with >10x coverage. Previously described haplogroups, such as European, Near Eastern, East Asian, and a haplogroup including the mtDNA sequences from Island Southeast Asian (ISEA) endemic *Sus* species: *S. celebensis* (Sulawesi), *S. cebifrons* (Philippines), *S. barbatus* (Borneo and Sumatra) and *S. verrucosus* (Java; Fig. S1), all formed well-supported clades. We also identified two well-supported clades corresponding to the Pacific Clade and the Lanyu Clade, both of which had previously been identified using short-fragment PCR data (*6*).

To quantify the frequency of both Pacific Clade and Lanyu Clade across ISEA and the Pacific, we combined this data with 703 control region fragment sequences including historical and ancient specimens, (*6, 13*), and a recently published dataset of 356 modern individuals from the Philippines (*20, 21*). We identified 58 individuals that possessed mitochondrial sequences belonging to the Lanyu haplogroup, all of which were found in the Philippines (45), Palawan (7) and Taiwan (6). Of the 90 individuals that possessed a Pacific Clade signature (Fig. 1B), the majority (55) were located east of the Wallace Line in Wallacea, New Guinea, Melanesia, and Polynesia. The remaining individuals were found across a geographically diverse range including Bali, Sumatra, Thailand, Laos, Japan, Yunnan, Nepal, and in a single modern individual from the Philippines. Of the 72 individual pigs from east of the Wallace line included in these data, ∼77% possessed haplotypes belonging to the Pacific clade, while ∼22% (16/72) possessed haplotypes belonging to the East Asian haplogroup and one individual from New Caledonia possessed a haplotype belonging to the European clade. These results indicate that, although commonly found east of the Wallace Line, the Pacific Clade haplogroup is not fixed in pigs from this region. Instead, it is found in pigs across East and South Asia, and in a low frequency in the Philippines (1/352).

These results suggest that the distinctive tooth shape (Pacific shape) previously associated with pigs carrying a Pacific Clade mtDNA signature may also be present west of the Wallace Line and in the Philippines. To address this, we conducted a morphometric analysis of pig teeth (third lower molar) based on 307 archaeological and 401 modern specimens. We used 12 landmarks and 87 sliding semilandmarks to capture tooth shape variation. This analysis identified a distinct tooth morphology (the Pacific shape) that is distinct from the tooth shape observed in archaeological pigs from mainland East Asia (correct cross-validation = 97% (CI: 92.3-100%). The Pacific shape was identified in pigs east of the Wallace Line including modern and ancient individuals from PNG, Vanuatu and the Marquesas (Fig. 1C).

Interestingly, none of the 36 archaeological individuals from the Philippines possessed the Pacific shape, while 3 out of 35 individuals from Taiwan were identified as having this shape with high probability (>92%). Five out of 14 individuals from Sarawak (Borneo) were also found to possess the Pacific shape. These results demonstrate that multiple pig populations found west of the Wallace linein Taiwan, the Philippines, Borneo, and mainland Asia carry these distinctive mtDNA and tooth shape signatures This geographic distribution makes it difficult to use these markers to establish the geographic origins of pigs found east of the Wallace Line, and our results suggest that these signatures may have become fixed through founder events as pig populations were transported into ISEA and Oceania..

### The ancestry of pigs east of the Wallace Line

To further address the origin of pigs east of the Wallace Line, we analysed 576 nuclear genomes with >0.1x coverage. We first conducted a Principal Component Analysis (PCA), that corroborated the existence of three major ancestry clines (Fig. 1D) observed in previous publications (*18, 22, 23*). Principal Component 1 (PC1) differentiated *S. scrofa* (wild and domestic) and the ISEA-endemic *Sus* species: *S. celebensis, S. cebifrons, S. philippensis, S. barbatus* and *S. verrucosus* (Fig. 1D). PC2 differentiated European and Asian *S. scrofa* populations (Fig. 1D). *S. scrofa* individuals from ISEA (including Wallacea), Micronesia, Melanesia, and Polynesia (represented by red triangles in Fig. 1A&D) are broadly distributed across PC1 and PC2 indicating that they possess mixed ancestry that includes ISEA-endemic *Sus*, Asian *S. scrofa* and European *S. scrofa*. These results are consistent with the observation that the majority of *S. scrofa* individuals from Sumatra, the Malay Peninsula and nearby islands also possessed ISEA-endemic *Sus* mtDNA haplogroups (Fig. 1B; Fig. S1).

To quantify the proportion of ancestry derived from ISEA-endemic *Sus*, Asian *S. scrofa* and European *S. scrofa* proportions in individual genomes, we used a combination of analyses including ADMIXTURE and D-statistics. We first employed an unsupervised ADMIXTURE analysis (K=3) based on 50 wild *S. scrofa* individuals from Europe and 40 from East Asia (China and Korea), in addition to the 30 ISEA-endemic *Sus* genomes in our dataset. This facilitated the identification of representative genomes for each ancestry (i.e. those that possessed >99% from only one source; Fig. S4). These representatives served as sources in a supervised ADMIXTURE (K=3) analysis that included the 576 genomes with >0.1x coverage. We further decomposed ISEA-endemic *Sus* ancestry into one of four components (*S. celebensis, S. cebifrons/S. philippensis, S. barbatus*, and *S. verrucosus*) using D-statistics (see Supplementary Materials).

These analyses confirmed the widespread presence of both ISEA-endemic *Sus* and Asian *S. scrofa* nuclear ancestry in individuals from ISEA that had been previously identified as a subspecies of *S. scrofa* based on morphological descriptions (Data S1). Many of these samples also possessed ISEA-endemic *Sus* mtDNA haplogroups, particularly in Wallacea, Sumatra, Java, and the Malay Peninsula (Fig. 1B; Fig. 2). There was also a strong concordance between geographical location and ISEA-endemic *Sus* ancestry: 18 *S. scrofa* individuals from Java and Sumatra possessed between ∼13-37% ancestry derived from either *S. verrucosus* (endemic to Java) or *S. barbatus* (endemic to Sumatra and Borneo), while 8 admixed individuals east of the Wallace Line possessed between ∼15-80% *S. celebensis* ancestry (endemic to Sulawesi; Fig. 2). Interestingly, we found little to no *S. cebifrons/S. philippensis* ancestry (endemic to the Philippines) in 14 *S. scrofa* individuals from the Philippines (Fig. 2).

**Fig. 2.**
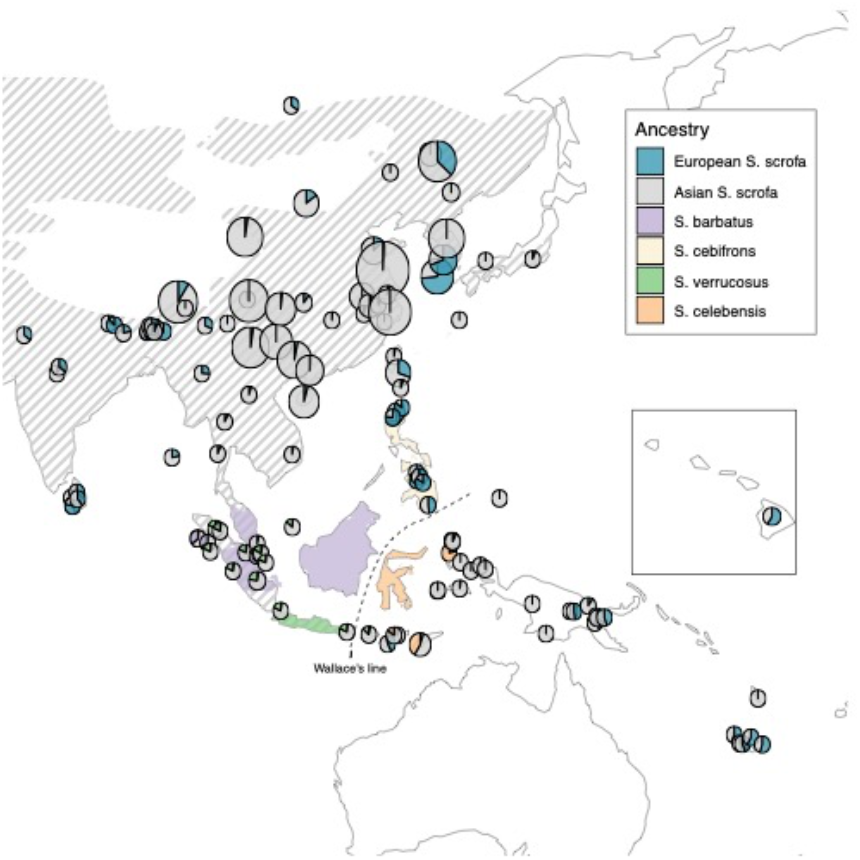
Ancestry of pigs inferred using ADMIXTURE. Results of a supervised ADMIXTURE analysis with K=3. Each pie chart represents a location, size is correlated with the number of samples.

Most individuals from Western Sunda (Sumatra, Java and nearby islands) possessed similar levels of *S. verrucosus* ancestry (13-18%). This finding supports previous work indicating that wild *S. scrofa* from Sumatra interbred with *S. verrucosus* during the Pleistocene when low sea levels allowed for the connection of Sumatra, Borneo, and Java via the Sunda Shelf, which facilitated migration and population exchange (*18, 19*). Three individuals from Simeulue Island, northwest of Sumatra, which were suspected to have been part of a population translocated from Sulawesi based on their morphological classification as *S. celebensis* (*24*), possessed ∼30% *S. barbatus* and ∼70% Asian *S. scrofa* ancestry. These results suggest that interbreeding between species can confuse morphological analyses and lead to spurious identifications.

Although all 12 individuals in Wallacea, outside of the natural range of *Sus* species (i.e. in Maluku and Lower Sunda) possessed some degree of Asian *S. scrofa* and/or *S. celebensis* ancestry, the proportion of each varied widely between islands. In the Maluku archipelago northeast of Sulawesi, *S. celebensis* ancestry ranged from 80% on Halmahera to ∼5% on Morotai, just a few kilometers north of Halmahera. Additionally, this ancestry was absent in the Central Maluku islands of Buru or Seram. The absence of *S. celebensis* ancestry within pigs found on Buru is puzzling given the presence of babirusa (*Babyrousa babyrussa*) on the island since this population has been thought to have been introduced from Sulawesi, and previous surveys have suggested that *S. celebensis* individuals also occurred on Buru (*25*).

A similar pattern was observed in the Lesser Sunda region of Wallacea including the islands of Timor, where *S. celebensis* ancestry ranged from 100% to 29%, and Flores, where it ranged from 16% to 5%. These results demonstrate that *S. celebensis* populations occur well beyond the native range of the species on Sulawesi. While natural dispersal to some islands cannot be ruled out, the open water distance between Sulawesi and Flores suggests that hunter-gatherers deliberately translocated this species during or since the Late Pleistocene (*6, 7*).

We did not detect any ISEA-endemic *Sus* ancestry in domestic pigs from the Philippines, Melanesia, and Polynesia. European ancestry was also absent from historic individuals from New Guinea dated to the early-mid to 20th century, but ranged from 10 to 58% in modern domestic New Guinea populations. In modern New Caledonian pigs, European ancestry ranged from ∼43-57%, and a pig from Hawaii, sampled in 1937, possessed 58% European ancestry. All recent domestic pigs from the Philippines possessed between 27-90% European ancestry. We also detected ∼42% European ancestry in Sumba, an island south of Flores.

The patterns of European and Asian *S. scrofa* ancestry covariance across the genome based on DATES (*26*) indicated that European *S. scrofa* ancestry was introduced to these islands during the 19th and early 20th centuries (Supplementary Materials). In addition, ancient genomes from Vanuatu (∼2,500 BP), the Philippines (∼500 BP), and Palau (∼700 BP) showed no evidence of European ancestry. These results demonstrate that European *S. scrofa* ancestry was introduced to these regions during the colonial period and has since risen to substantial proportions in pig populations across the region.

### The geographic origin of Asian S. scrofa ancestry east of the Wallace Line

Although multiple historical and modern individuals east of the Wallace Line possess a diverse mix of ancestry, including *S. celebensis* and European *S. scrofa*, they all possess some degree of Asian *S. scrofa* ancestry. Assessing the origin of this Asian *S. scrofa* ancestry is challenging, however, particularly in mixed genomes that also feature European *S. scrofa* or ISEA-endemic *Sus* ancestry. In addition, the Asian *S. scrofa* lineage is widespread and encompasseswild pigs from North and South China, Korea, Japan, ISEA (Sumatra and Java), ancient pigs from Melanesia (∼2,900 BP), the Philippines and Micronesia (Palau), and East Asian domestic pigs.

To overcome these constraints,, we performed ancestry deconvolution, a process by which specific ancestries in admixed individuals can be isolated using local ancestry inference. We first phased all genomes with >8x coverage using SHAPEIT v4 (*27*), and imputed the genomes with >1x coverage using GLIMPSE (*28*). We then used GNOMIX (*29*) to identify ancestry blocks from Asian *S. scrofa*, European *S. scrofa*, and ISEA-endemic *Sus* ancestry, using wild, un-admixed, individuals as reference (see Supplementary Materials). We first assessed how imputation of lower-coverage genomes influenced our ability to conduct local ancestry inference by downsampling high-coverage individuals. This analysis showed that GNOMIX local ancestry inference is highly accurate (94-99%) even when applied to individuals with coverage as low as 0.1x (Fig. S3).

For each admixed genome, we filtered ancestry blocks assigned to European *S. scrofa* and ISEA-endemic *Sus* ancestry. We then computed f4 ratios to compare levels of non-Asian *S. scrofa* ancestry before and after filtering out both European *S. scrofa* and ISEA-endemic *Sus* ancestry blocks (Fig. 3A). European *S. scrofa* ancestry in admixed individuals from New Caledonia, Hawaii, Papua New Guinea and Taiwan (Lanyu pigs) decreased from ∼25-50% to 0-5%, while European *S. scrofa* ancestry in domestic pigs from the Philippines decreased from ∼30-70% to 0-15%. ISEA-endemic *Sus* (*S. celebensis*) ancestry in individuals from Wallacea, including Flores, Timor, and Halmahera, decreased from between ∼80-40% to below 5%.

**Fig. 3.**
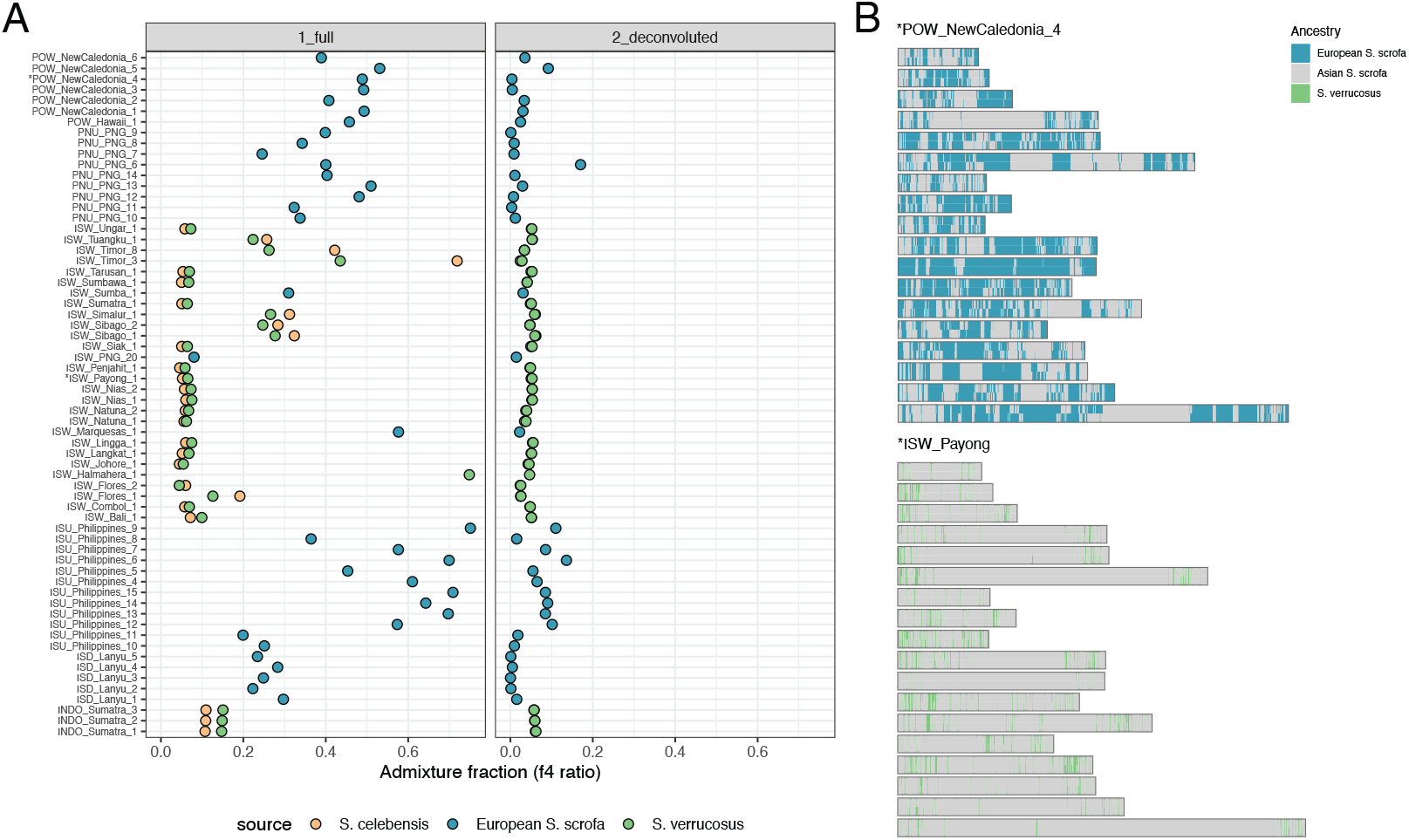
(A) Accuracy of ancestry deconvolution. European S. scrofa / ISEA-endemic Sus ancestry proportion in genomes of mixed individuals before (full) and after (deconvoluted) ancestry deconvolution **(B). Size of ancestry blocks in two genomes**. Blocks of ancestry across pig chromosomes from a New Caledonian feral individual and a wild pig from Payong.

ISEA-endemic *Sus* (*S. verrucosus*) ancestry in wild pigs from Western Indonesia, in Java, Sumatra, and nearby islands, however, decreased from between 5-14% to ∼5%. This lower deconvolution success in these individuals can be explained by ancient admixture that took place during the Pleistocene (*18, 19*). This ancient gene flow left small blocks of ISEA-endemic *Sus* ancestry across the genome that GNOMIX finds more difficult to identify. In contrast, due to its recent introduction in the region, the European *S. scrofa* ancestry is present in larger blocks (Fig. 3B), and is thus more easily identified. The wild pigs from Western Indonesia could also possess *S. scrofa* ancestry that is not well characterized in our panels.

To assess whether Western Indonesian *S. scrofa* possesses ancestry that is not represented by either Asian (represented by Chinese wild pigs) or European *S. scrofa* (represented by European wild boar), we fitted a range of ancestry models using Admixtools2. Specifically, for each ISEA individual in our dataset that possessed *S. scrofa* ancestry, we fitted models where its ancestry is derived either from Asian *S. scrofa*, or from an unsampled sister-lineage equally distant to Asian and European *S. scrofa* (Fig. S5). Models with ancestry from the sister-lineage to Asian and European *S. scrofa* tended to fit better for Western Indonesian *S. scrofa*, while models with Asian *S. scrofa* ancestry fitted better for populations east of the Wallace Line (Fig. S5). This suggests that Western Indonesian *S. scrofa* possesses ancestry from a population that is equally distant from Asian and European *S. scrofa*. Our findings corroborate previous studies that have suggested that Western Indonesian *S. scrofa* represent an early offshoot that diverged from other mainland Eurasian *S. scrofa* populations during the Pleistocene (*18*).

We next assessed the region of origin of Asian *S. scrofa* ancestry in Wallacean, Melanesian, Micronesian and Polynesian pigs. To do so, we used outgroup-*f3* to calculate shared drift between a ∼100-year-old genome from a Papuan feral pig (ISW_PNG_18) and all other genomes, without European *S. scrofa* and ISEA-endemic ancestry (Fig. 3A). We chose this Papuan individual because its ancestry is similar to that of a low-coverage (0.7x) ancient individual from Vanuatu (∼2,900 BP; Supplementary Materials) excavated from an Austronesian-associated Lapita culture archeological site. This analysis demonstrated that all *S. scrofa* individuals from Wallacea, Micronesia, Melanesia, and Polynesia share much stronger affinities with ISW_PNG_18, than with *S. scrofa* from Western Indonesia (Sumatra and Java).

This analysis also indicated that ISW_PNG_18 shares more drift with the Asian *S. scrofa* ancestry component found in other pigs east of the Wallace line, from Maluku to Hawaii (Fig. 4A). West and North of the Wallace line, ISW_PNG_18 shared more drift with ISU_UchuPalau_,1 a 672-year-old Micronesian pig from Palau and ISU_Philippines_1; a 500-year-old individual from the Batanes Islands. ISW_PNG_18 also shared more drift with domestic pigs from Taiwan (Lanyu pigs), and South China (e.g., Bamaxiang, Luchuan and Wuzhishan pigs from Hainan Island and adjacent mainland Chinese provinces) than with domestic pigs from Northern China and Vietnam, or with free-living Western Indonesian (Sumatra and Java) and wild Chinese *S. scrofa* (Fig. 4A).

**Fig. 4.**
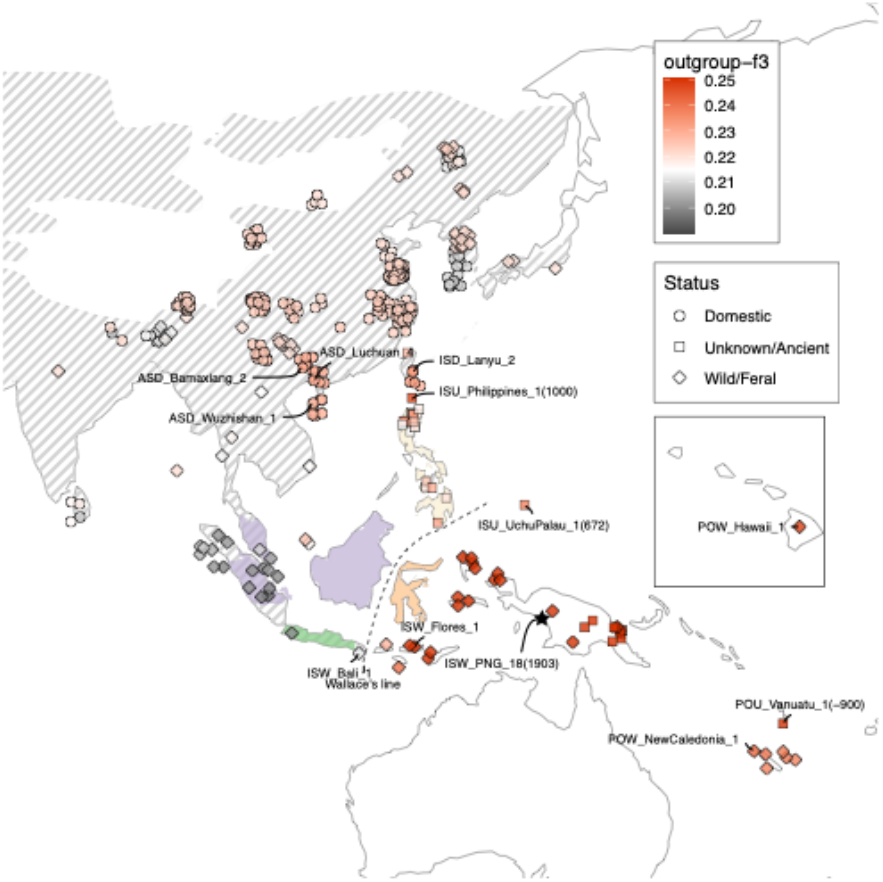
Outgroup-f3 after deconvolution. Outgroup-f3 (shared drift) between genomes plotted on the map and ISW_PNG_18 (feral pig from New Guinea) after removing European and ISEA endemic-*Sus* ancestry from the genome of mixed individuals using ancestry deconvolution. Higher outgroup-f3 values indicate higher degree of genetic affinity with the ISW_PNG_18 individual which is used as a representative of Austronesian pig ancestry.

Combined, these results suggest that the domestic and feral pigs found today beyond the natural range of *S. scrofa* in Wallacea and the Pacific, are closer genetically to ancient pigs in the Philippines, and modern domestic pigs in Taiwan, than to modern and historical individuals from peninsular Southeast Asia and Western Indonesia. This indicates that pigs in the Pacific are, at least partly, descendants of domestic pigs introduced to the region via the Philippines, most probably by Austronesian-speaking groups who possessed East Asian human ancestry (*17*).

### Multiple waves of dispersal and introduction in the Lesser Sunda Islands

Interestingly, a sample from Sumbawa, a few hundred kilometres east of the Wallace Line, possesses a mixture of Asian *S. scrofa* and *S. verrucosus* ancestry (endemic to Java), similar to pigs in Bali. Pigs from Flores and Timor (Fig. 2), however, instead possess a mixture of Asian *S. scrofa* and *S. celebensis* ancestry (endemic to Sulawesi). Admixture modeling further suggests that at least part of the *S. scrofa* component in the Sumbawa pig is derived from a sister-lineage to Asian *S. scrofa* and European *S. scrofa*, similar to that observed in Western Indonesian pigs (Fig. S5). These results suggest that Sumbawa pigs may, to some extent, possess ancestry from the western Lesser Sunda Islands, suggesting a possible natural and/or human-mediated translocation of pigs across the Wallace Line in the Lesser Sunda Islands. This ancestry pattern in the Lesser Sunda Islands illustrates the complex evolutionary history of pigs east of the Wallace Line.

Combined, our results suggest multiple dispersal processes in the region that possibly started with a natural and/or human-mediated translocation of Western Indonesian pigs from Bali across the Wallace Line that possessed Western Indonesian *S. scrofa* and *S. verrucosus* ancestry. This was likely followed by the introduction of *S. celebensis* from Sulawesi, potentially by hunter-gatherer populations. Subsequently, domestic *S. scrofa* pigs derived from populations in mainland Asia were introduced into the region from northern Wallacea (following their introduction from Southern China and the Philippines) sometime after 3,500-3,000 years ago. Many of these introduced pigs later became feral on islands across the region. Finally, European pigs were introduced broadly across the region, including onto the Lesser Sundas to the island of Sumba during the colonial period.

### Conclusions

The natural and anthropogenic history of pigs east of the Wallace Line is complex. We found that most free-living and domestic pigs across a vast geographical range from the Philippines to Hawaii possess a degree of ancestry linked to domestic pigs (*S. scrofa*) likely introduced to ISEA during the spread of Austronesian-speaking groups from Southeast China and Taiwan via the Philippines ∼4,000 years ago. Interestingly, pigs in Melanesia, Micronesia, and Polynesia lack any detectable ancestry from wild endemic *Sus* species present along the Austronesian dispersal route in the Philippines, Sulawesi and other islands. This suggests that the first wave of introduced pigs were isolated from local, indigenous *Sus* populations. This lack of gene flow mirrors the pattern observed in the human populations who transported the pigs, and who did not interbreed with local groups during the initial phases of their dispersal to Melanesia (*17*). In contrast to human populations, where mixing occurred between incoming Austronesian groups and local Papuan groups a few hundred years after their arrival in Melanesia (*15, 16*), no native *Sus* species existed in Melanesia at the time of the introduction of domestic pigs. This prevented interbreeding between the introduced pigs and any indigenous pig populations. Only those populations left behind in Wallacea, which subsequently became feral, had the chance to hybridize with indigenous wild species, such as *Sus celebensis*.

The deliberate human-mediated dispersal of *S. scrofa* from mainland Asia to Oceania, and their initial isolation from local wild species may reflect the fact that these introduced pigs possessed specific domestic traits that facilitated their transport and management by Austronesian-speaking groups. Transporting these animals across islands resulted in a unique demographic history characterized by serial founder effects, potentially explaining observed patterns in mitochondrial DNA and tooth morphology, that has since left lasting genomic signatures. Future research utilizing high-coverage genomes from modern, historical, and archaeological samples will be crucial to unravel the demographic history of these populations.

## Acknowledgments

HPC computing was performed on the BioHPC (DFG INST 86/2050-1 FUGG) and Linux-Cluster of the Leibniz Supercomputing Centre (LRZ Munich).

## Author contributions

Conceptualization: LAFF, GL, KD

Data acquisition: AL, AE, AM, CL, OL, RD, KJM, RB

Data analysis: DWGS, AM, AE, KT, LAFF

Sample acquisition: AA, PB, SB, PB, AB, TCT, GC, RC, TC, MSE, LGF, PG, MG, BH, SH, HH, KMH, MJH, TH, ACK, CL, AAM, H-JM, EJ, CM, KM, KN, MO, RP PP, KS, LS, PS, MS, ST, USV, JG, TD

Funding acquisition: LAFF, GL, KD Writing – original draft: LAFF, DWGS

Writing – review & editing: GL, RD, KM, TD, MPC, AK, KMH, MS

## Data and materials availability

The reads for the data have been deposited at the European Nucleotide Archive (ENA) with project number PRJEB83975.

## Supplementary Materials

Materials and Methods

Figs. S1 to S9

Data S1 to S3

References (*30*–*58*)

## References and Notes

1. D. R. Spatz, K. M. Zilliacus, N. D. Holmes, S. H. M. Butchart, P. Genovesi, G. Ceballos, B. R. Tershy, D. A. Croll, Globally threatened vertebrates on islands with invasive species. Sci Adv 3, e1603080 (2017).

2. M. J. Struebig, S. G. Aninta, M. Beger, A. Bani, H. Barus, S. Brace, Z. G. Davies, M. D. Brauwer, K. Diele, C. Djakiman, R. Djamaluddin, R. Drinkwater, A. Dumbrell, D. Evans, M. Fusi, L. Herrera-Alsina, D. T. Iskandar, J. Jompa, B. Juliandi, L. T. Lancaster, sG. Limmon, Lindawati, M. G. Y. Lo, P. Lupiyaningdyah, M. McCannon, E. Meijaard, S. L. Mitchell, S. Mumbunan, D. O’Connell, O. G. Osborne, A. S. T. Papadopulos, J. S. Rahajoe Rosaria,, S. J. Rossiter Rugayah,, H. Rustiami, U. Salzmann Sheherazade,, I. M. Sudiana, E. Sukara, J. S. Tasirin, A. Tjoa, J. M. J. Travis, L. Trethowan, A. Trianto, T. Utteridge, M. Voigt, N. Winarni, Z. Zakaria, D. P. Edwards, L. Frantz, J. Supriatna, Safeguarding Imperiled Biodiversity and Evolutionary Processes in the Wallacea Center of Endemism. Bioscience 72, 1118–1130 (2022).

3. T. Heinsohn, Animal translocation: long-term human influences on the vertebrate zoogeography of Australasia (natural dispersal versus ethnophoresy). Aust. Zool. 32, 351–376 (2003).

4. A. A. Oktaviana, R. Joannes-Boyau, B. Hakim, B. Burhan, R. Sardi, S. Adhityatama, Hamrullah, I. Sumantri, M. Tang, R. Lebe, I. Ilyas, A. Abbas, A. Jusdi, D. E. Mahardian, S. Noerwidi, M. N. R. Ririmasse, I. Mahmud, A. Duli, L. M. Aksa, D. McGahan, P. Setiawan, A. Brumm, M. Aubert, Narrative cave art in Indonesia by 51,200 years ago. Nature 631, 814–818 (2024).

5. M. Aubert, A. Brumm, M. Ramli, T. Sutikna, E. W. Saptomo, B. Hakim, M. J. Morwood, G. D. van den Bergh, L. Kinsley, A. Dosseto, Pleistocene cave art from Sulawesi, Indonesia. Nature 514, 223–227 (2014).

6. G. Larson, T. Cucchi, M. Fujita, E. Matisoo-Smith, J. Robins, A. Anderson, B. Rolett, M. Spriggs, G. Dolman, T.-H. Kim, N. T. D. Thuy, E. Randi, M. Doherty, R. A. Due, R. Bollt, T. Djubiantono, B. Griffin, M. Intoh, E. Keane, P. Kirch, K.-T. Li, M. Morwood, L. M. Pedriña, P. J. Piper, R. J. Rabett, P. Shooter, G. Van den Bergh, E. West, S. Wickler, J. Yuan, A. Cooper, K. Dobney, Phylogeny and ancient DNA of Sus provides insights into neolithic expansion in Island Southeast Asia and Oceania. Proc. Natl. Acad. Sci. U. S. A. 104, 4834–4839 (2007).

7. G. D. van den Bergh, H. J. M. Meijer, R. Due Awe, M. J. Morwood, K. Szabó, L. W. van den Hoek Ostende, T. Sutikna, E. W. Saptomo, P. J. Piper, K. M. Dobney, The Liang Bua faunal remains: a 95k.yr. sequence from Flores, East Indonesia. J. Hum. Evol. 57, 527– 537 (2009).

8. T. Sutikna, M. W. Tocheri, J. T. Faith, Jatmiko, R. Due Awe, H. J. M. Meijer, E. Wahyu Saptomo, R. G. Roberts, The spatio-temporal distribution of archaeological and faunal finds at Liang Bua (Flores, Indonesia) in light of the revised chronology for Homo floresiensis. J. Hum. Evol. 124, 52–74 (2018).

9. P. Bellwood, First Farmers (Wiley-Blackwell, Hoboken, NJ, 2023).

10. F. Petchey, M. Spriggs, S. Bedford, F. Valentin, The chronology of occupation at Teouma, Vanuatu: Use of a modified chronometric hygiene protocol and Bayesian modeling to evaluate midden remains. J. Archaeol. Sci. Rep. 4, 95–105 (2015).

11. Lapita in the Southwest Pacific: Origins, Distribution, Chronology, Economy, and Transformation.

12. F. Valentin, H. R. Buckley, E. Herrscher, R. Kinaston, S. Bedford, M. Spriggs, S. Hawkins, K. Neal, Lapita subsistence strategies and food consumption patterns in the community of Teouma (Efate, Vanuatu). J. Archaeol. Sci. 37, 1820–1829 (2010).

13. G. Larson, R. Liu, X. Zhao, J. Yuan, D. Fuller, L. Barton, K. Dobney, Q. Fan, Z. Gu, X.-H. Liu, Y. Luo, P. Lv, L. Andersson, N. Li, Patterns of East Asian pig domestication, migration, and turnover revealed by modern and ancient DNA. Proc. Natl. Acad. Sci. U. S. A. 107, 7686–7691 (2010).

14. T. Denham, Early farming in Island Southeast Asia: an alternative hypothesis. Antiquity 87, 250–257 (2013).

15. M. Lipson, P. Skoglund, M. Spriggs, F. Valentin, S. Bedford, R. Shing, H. Buckley, I. Phillip, G. K. Ward, S. Mallick, N. Rohland, N. Broomandkhoshbacht, O. Cheronet, M. Ferry, T. K. Harper, M. Michel, J. Oppenheimer, K. Sirak, K. Stewardson, K. Auckland, A. V. S. Hill, K. Maitland, S. J. Oppenheimer, T. Parks, K. Robson, T. N. Williams, D. J. Kennett, A. J. Mentzer, R. Pinhasi, D. Reich, Population turnover in Remote Oceania shortly after initial settlement. Curr. Biol. 28, 1157–1165.e7 (2018).

16. C. Posth, K. Nägele, H. Colleran, F. Valentin, S. Bedford, K. W. Kami, R. Shing, H. Buckley, R. Kinaston, M. Walworth, G. R. Clark, C. Reepmeyer, J. Flexner, T. Maric, J. Moser, J. Gresky, L. Kiko, K. J. Robson, K. Auckland, S. J. Oppenheimer, A. V. S. Hill, A. J. Mentzer, J. Zech, F. Petchey, P. Roberts, C. Jeong, R. D. Gray, J. Krause, A. Powell, Language continuity despite population replacement in Remote Oceania. Nat. Ecol. Evol. 2, 731–740 (2018).

17. P. Skoglund, C. Posth, K. Sirak, M. Spriggs, F. Valentin, S. Bedford, G. R. Clark, C. Reepmeyer, F. Petchey, D. Fernandes, Q. Fu, E. Harney, M. Lipson, S. Mallick, M. Novak, N. Rohland, K. Stewardson, S. Abdullah, M. P. Cox, F. R. Friedlaender, J. S. Friedlaender, T. Kivisild, G. Koki, P. Kusuma, D. A. Merriwether, F.-X. Ricaut, J. T. S. Wee, N. Patterson, J. Krause, R. Pinhasi, D. Reich, Genomic insights into the peopling of the Southwest Pacific. Nature 538, 510–513 (2016).

18. L. A. F. Frantz, J. G. Schraiber, O. Madsen, H.-J. Megens, M. Bosse, Y. Paudel, G. Semiadi, E. Meijaard, N. Li, R. P. M. A. Crooijmans, A. L. Archibald, M. Slatkin, L. B. Schook, G. Larson, M. A. M. Groenen, Genome sequencing reveals fine scale diversification and reticulation history during speciation in Sus. Genome Biol. 14, R107 (2013).

19. L. A. F. Frantz, O. Madsen, H.-J. Megens, M. A. M. Groenen, K. Lohse, Testing models of speciation from genome sequences: divergence and asymmetric admixture in Island South-East Asian Sus species during the Plio-Pleistocene climatic fluctuations. Mol. Ecol. 23, 5566–5574 (2014).

20. J. K. N. Layos, C. J. P. Godinez, L. M. Liao, Y. Yamamoto, J. S. Masangkay, H. Mannen, M. Nishibori, Origin and demographic history of Philippine pigs inferred from mitochondrial DNA. Front. Genet. 12 (2022).

21. J. B. Banayo, K. L. V. Manese, A. J. Salces, T. Yamagata, Phylogeny and genetic diversity of Philippine native pigs (Sus scrofa) as revealed by mitochondrial DNA analysis. Biochem. Genet. 61, 1401–1417 (2023).

22. L. A. F. Frantz, J. G. Schraiber, O. Madsen, H.-J. Megens, A. Cagan, M. Bosse, Y. Paudel, R. P. M. A. Crooijmans, G. Larson, M. A. M. Groenen, Evidence of long-term gene flow and selection during domestication from analyses of Eurasian wild and domestic pig genomes. Nat. Genet. 47, 1141–1148 (2015).

23. L. A. F. Frantz, J. Haile, A. T. Lin, A. Scheu, C. Geörg, N. Benecke, M. Alexander, A. Linderholm, V. E. Mullin, K. G. Daly, V. M. Battista, M. Price, K. J. Gron, P. Alexandri, R.-M. Arbogast, B. Arbuckle, A. Bӑlӑşescu, R. Barnett, L. Bartosiewicz, G. Baryshnikov, C. Bonsall, D. Borić, A. Boroneanţ, J. Bulatović, C. Çakirlar, J.-M. Carretero, J. Chapman, M. Church, R. Crooijmans, B. D. Cupere, C. Detry, V. Dimitrijevic, V. Dumitraşcu, L. du Plessis, C. J. Edwards, C. M. Erek, A. Erim-Özdoğan, A. Ervynck, D. Fulgione, M. Gligor, A. Götherström, L. Gourichon, M. A. M. Groenen, D. Helmer, H. Hongo, L. K. Horwitz, E. K. Irving-Pease, O. Lebrasseur, J. Lesur, C. Malone, N. Manaseryan, A. Marciniak, H. Martlew, M. Mashkour, R. Matthews, G. M. Matuzeviciute, S. Maziar, E. Meijaard, T. McGovern, H.-J. Megens, R. Miller, A. F. Mohaseb, J. Orschiedt, D. Orton, A. Papathanasiou, M. P. Pearson, R. Pinhasi, D. Radmanović, F.-X. Ricaut, M. Richards, R. Sabin, L. Sarti, W. Schier, S. Sheikhi, E. Stephan, J. R. Stewart, S. Stoddart, A. Tagliacozzo, N. Tasić, K. Trantalidou, A. Tresset, C. Valdiosera, Y. van den Hurk, S. Van Poucke, J.-D. Vigne, A. Yanevich, A. Zeeb-Lanz, A. Triantafyllidis, M. T. P. Gilbert, J. Schibler, P. Rowley-Conwy, M. Zeder, J. Peters, T. Cucchi, D. G. Bradley, K. Dobney, J. Burger, A. Evin, L. Girdland-Flink, G. Larson, Ancient pigs reveal a near-complete genomic turnover following their introduction to Europe. Proceedings of the National Academy of Sciences 116, 17231– 17238 (2019).

24. C. P. Groves, Ancestors for the Pigs : Taxonomy and Phylogeny of the Genus Sus (Australian National University Press, Canberra, 1981).

25. IUCN, Sus celebensis: Burton, J., Mustari, A. & Rejeki, I, IUCN (2016); 10.2305/iucn.uk.2020-2.rlts.t41773a44141588.en.

26. M. Chintalapati, N. Patterson, P. Moorjani, The spatiotemporal patterns of major human admixture events during the European Holocene. Elife 11, e77625 (2022).

27. O. Delaneau, J.-F. Zagury, M. R. Robinson, J. L. Marchini, E. T. Dermitzakis, Accurate, scalable and integrative haplotype estimation. Nat. Commun. 10, 5436 (2019).

28. S. Rubinacci, D. M. Ribeiro, R. J. Hofmeister, O. Delaneau, Efficient phasing and imputation of low-coverage sequencing data using large reference panels. Nat. Genet. 53, 120–126 (2021).

29. H. Hilmarsson, A. S. Kumar, R. Rastogi, C. D. Bustamante, D. M. Montserrat, A. G. Ioannidis, High resolution ancestry deconvolution for next generation genomic data, bioRxiv (2021). 10.1101/2021.09.19.460980.

30. B. Egloff, Recent prehistory in southeast Papua. (1979).

31. W. B. Masse, “The archaeology and ecology of fishing in the Belau Islands, Micronesia,” thesis, Southern Illinois University (1989).

32. G. Clark, C. Reepmeyer, Last millennium climate change in the occupation and abandonment of Palau’s Rock Islands. Archaeol. Ocean. 47, 29–38 (2012).

33. P. Bellwood, E. Dizon, “The batanes islands, their first observers, and previous archaeology” in 4000 Years of Migration and Cultural Exchange (Terra Australis 40): The Archaeology of the Batanes Islands, Northern Philippines (ANU Press, 2013), pp. 1– 8.

34. P. J. Reimer, W. E. N. Austin, E. Bard, A. Bayliss, P. G. Blackwell, C. Bronk Ramsey, M. Butzin, H. Cheng, R. L. Edwards, M. Friedrich, P. M. Grootes, T. P. Guilderson, I. Hajdas, T. J. Heaton, A. G. Hogg, K. A. Hughen, B. Kromer, S. W. Manning, R. Muscheler, J. G. Palmer, C. Pearson, J. van der Plicht, R. W. Reimer, D. A. Richards, E. M. Scott, J. R. Southon, C. S. M. Turney, L. Wacker, F. Adolphi, U. Büntgen, M. Capano, S. M. Fahrni, A. Fogtmann-Schulz, R. Friedrich, P. Köhler, S. Kudsk, F. Miyake, J. Olsen, F. Reinig, M. Sakamoto, A. Sookdeo, S. Talamo, The IntCal20 Northern hemisphere radiocarbon age calibration curve (0–55 cal kBP). Radiocarbon 62, 725–757 (2020).

35. C. Ottoni, L. G. Flink, A. Evin, C. Geörg, B. De Cupere, W. Van Neer, L. Bartosiewicz, A. Linderholm, R. Barnett, J. Peters, R. Decorte, M. Waelkens, N. Vanderheyden, F.-X. Ricaut, C. Cakirlar, O. Cevik, A. R. Hoelzel, M. Mashkour, A. F. M. Karimlu, S. S. Seno, J. Daujat, F. Brock, R. Pinhasi, H. Hongo, M. Perez-Enciso, M. Rasmussen, L. Frantz, H.-J. Megens, R. Crooijmans, M. Groenen, B. Arbuckle, N. Benecke, U. S. Vidarsdottir, J. Burger, T. Cucchi, K. Dobney, G. Larson, Pig domestication and human-mediated dispersal in western Eurasia revealed through ancient DNA and geometric morphometrics. Mol. Biol. Evol. 30, 824–832 (2013).

36. P. B. Damgaard, A. Margaryan, H. Schroeder, L. Orlando, E. Willerslev, M. E. Allentoft, Improving access to endogenous DNA in ancient bones and teeth. Sci. Rep. 5, 11184 (2015).

37. J. Dabney, M. Knapp, I. Glocke, M.-T. Gansauge, A. Weihmann, B. Nickel, C. Valdiosera, N. García, S. Pääbo, J.-L. Arsuaga, M. Meyer, Complete mitochondrial genome sequence of a Middle Pleistocene cave bear reconstructed from ultrashort DNA fragments. Proc. Natl. Acad. Sci. U. S. A. 110, 15758–15763 (2013).

38. C. Carøe, S. Gopalakrishnan, L. Vinner, S. S. T. Mak, M. H. S. Sinding, J. A. Samaniego, N. Wales, T. Sicheritz-Pontén, M. T. P. Gilbert, Single-tube library preparation for degraded DNA. Methods Ecol. Evol. 9, 410–419 (2018).

39. S. A. Miller, D. D. Dykes, H. F. Polesky, A simple salting out procedure for extracting DNA from human nucleated cells. Nucleic Acids Res. 16, 1215 (1988).

40. M. Meyer, M. Kircher, Illumina sequencing library preparation for highly multiplexed target capture and sequencing. Cold Spring Harb. Protoc. 2010, db.prot5448 (2010).

41. M.-T. Gansauge, M. Meyer, Single-stranded DNA library preparation for the sequencing of ancient or damaged DNA. Nat. Protoc. 8, 737–748 (2013).

42. S. Chen, Y. Zhou, Y. Chen, J. Gu, fastp: an ultra-fast all-in-one FASTQ preprocessor. Bioinformatics 34, i884–i890 (2018).

43. H. Li, R. Durbin, Fast and accurate short read alignment with Burrows-Wheeler transform. Bioinformatics 25, 1754–1760 (2009).

44. J. A. Fellows Yates, T. C. Lamnidis, M. Borry, A. Andrades Valtueña, Z. Fagernäs, S. Clayton, M. U. Garcia, J. Neukamm, A. Peltzer, Reproducible, portable, and efficient ancient genome reconstruction with nf-core/eager. PeerJ 9, e10947 (2021).

45. A. Stamatakis, RAxML-VI-HPC: maximum likelihood-based phylogenetic analyses with thousands of taxa and mixed models. Bioinformatics 22, 2688–2690 (2006).

46. K. Katoh, J. Rozewicki, K. D. Yamada, MAFFT online service: multiple sequence alignment, interactive sequence choice and visualization. Brief. Bioinform. 20, 1160– 1166 (2019).

47. L.-T. Nguyen, H. A. Schmidt, A. von Haeseler, B. Q. Minh, IQ-TREE: a fast and effective stochastic algorithm for estimating maximum-likelihood phylogenies. Mol. Biol. Evol. 32, 268–274 (2015).

48. S. Kalyaanamoorthy, B. Q. Minh, T. K. F. Wong, A. von Haeseler, L. S. Jermiin, ModelFinder: fast model selection for accurate phylogenetic estimates. Nat. Methods 14, 587–589 (2017).

49. D. T. Hoang, O. Chernomor, A. von Haeseler, B. Q. Minh, L. S. Vinh, UFBoot2: Improving the ultrafast bootstrap approximation. Mol. Biol. Evol. 35, 518–522 (2018).

50. M. Johnsson, A. Whalen, R. Ros-Freixedes, G. Gorjanc, C.-Y. Chen, W. O. Herring, D.-J. de Koning, J. M. Hickey, Genetic variation in recombination rate in the pig. Genet. Sel. Evol. 53, 54 (2021).

51. S. Purcell, B. Neale, K. Todd-Brown, L. Thomas, M. A. R. Ferreira, D. Bender, J. Maller, P. Sklar, P. I. W. de Bakker, M. J. Daly, P. C. Sham, PLINK: a tool set for whole-genome association and population-based linkage analyses. Am. J. Hum. Genet. 81, 559–575 (2007).

52. J. Meisner, S. Liu, M. Huang, A. Albrechtsen, Large-scale inference of population structure in presence of missingness using PCA. Bioinformatics 37, 1868–1875 (2021).

53. P. Librado, L. Orlando, Struct-f4: a Rcpp package for ancestry profile and population structure inference from f4-statistics. Bioinformatics 38, 2070–2071 (2022).

54. S. White, From Globalized Pig Breeds to Capitalist Pigs: A Study in Animal Cultures and Evolutionary History. Environ. Hist. Durh. N. C. 16, 94–120 (2011).

55. A. Evin, K. Dobney, R. Schafberg, J. Owen, U. S. Vidarsdottir, G. Larson, T. Cucchi, Phenotype and animal domestication: A study of dental variation between domestic, wild, captive, hybrid and insular Sus scrofa. BMC Evol. Biol. 15, 6 (2015).

56. F. J. Rohlf, tpsDig, version 2.10. (2006).

57. S. Schlager, “Chapter 9 - Morpho and Rvcg – Shape Analysis in R: R-Packages for Geometric Morphometrics, Shape Analysis and Surface Manipulations” in Statistical Shape and Deformation Analysis, G. Zheng, S. Li, G. Székely, Eds. (Academic Press, 2017), pp. 217–256.

58. A. Evin, L. G. Flink, A. Bălăşescu, D. Popovici, R. Andreescu, D. Bailey, P. Mirea, C. Lazăr, A. Boroneanţ, C. Bonsall, U. S. Vidarsdottir, S. Brehard, A. Tresset, T. Cucchi, G. Larson, K. Dobney, Unravelling the complexity of domestication: a case study using morphometrics and ancient DNA analyses of archaeological pigs from Romania. Philos. Trans. R. Soc. Lond. B Biol. Sci. 370, 20130616 (2015).

