## Supplementary materials for "Genomic and morphometric evidence for Austronesian-mediated pig translocation in the Pacific"

### Materials and Methods

Subhead

Type or paste text here. You can break this section up into subheads as needed (e.g., one on “Materials” and one on “Methods”).

Archeological specimens and site description of DNA samples

Uripiv, Vanuatu (2900-2000 BP; POU_Vanuatu_3; contact: Stuart Bedford)

The small islands of Vao and Uripiv are located off the northeast coast of Malekula Island in northern Vanuatu. Excavations on uplifted beach terraces on the sheltered coast of the islands revealed dense concentrations of midden deposits associated with the colonising Lapita phase (c. 2900 BP) through to c. 2000 BP. The sites are rich in faunal remains, both terrestrial and marine, shellfish, pottery and other artefacts. The sites on both islands were deeply stratified, up to 1.8 m deep to the sterile beach sand in some places, and generally very well preserved showing limited disturbance.

Teouma, Vanuatu (3000-2600 BP; POU_Vanuatu_4, POU_Vanuatu_5; contact: Stuart Bedford)

The Teouma Lapita site is located on the south coast of Efate Island, Vanuatu, and is currently 800 metres from Teouma Bay. Tectonic uplift, volcanic ashfall prior to and during the period of human utilisation of the site, and alluvial deposition from an adjacent stream have altered its immediate coastal location during the Lapita period at c. 3000 BP. Excavations by a joint ANU-Vanuatu National Museum team took place between 2004 and 2010, during which 68 burial features, including remains of just over 100 individuals were found. Teouma has numerous indicators of being an initial colonisation site for Efate, including extinct faunal remains, early ceramic forms and decoration, and exotic New Britain obsidian. This combined with the extensive and previously very rare Lapita skeletal remains underline its significance to those investigating colonisation in this region. There is evidence of continuing occupation at the site to c. 2600 BP. During the Lapita phase the site was composed of a cemetery and a contemporary settlement with a midden deposit. During later phases settlement expanded across the cemetery area.

Wanigela, Papua New Guinea (600-700 BP; PNU_PNG_2, PNU_PNG_3; contact: Geoffrey Clark)

The 14C sample was an adult pig tooth and fragment of mandible from Oreresan Village, Mound B (Spit 8 mixed) dated by Wk-30430, CRA 675 +/- 30, 13C -20.1, C:N 3.3. All the aDNA samples are from the Mound B excavation which is located in Oreresan Village located in coastal Eastern New Guinea in Collingwood Bay. The mound was excavated by Brian Egloff in the 1960s and site details are in his thesis publication (*30*).

Mound B is an oval-shaped mound near the Sasap River that is around 40 m in length and 17 m in width with a maximum height of 1.4 m. Excavations at the site totalled 22 m2 with sediments divided into eight zones and subzones – Zone I (top), Zone IIA, IIB, IIC, Zone III, Zone IVA, IVB (base). Pig bones were common in all excavation levels and comprised 80% of the identifiable vertebrate remains (*30*). Many pigs had been killed before their third molars had erupted when the body weight was estimated to be less than 80-120 kg. Four radiocarbon results obtained by Egloff and suggested that the Mound B materials were deposited between AD 100 and AD 1450. The large amount of pig remains in Mound B might represent debris from ceremonial feasting indicating that the importance of pigs in traditional societies in New Guinea dates to AD 1300-1400 (Calib 7.10 result for Wk-30430 at 95.4% is AD 1270-1390).

Uchul A Rois, Palau (900-1450 AD; ISU_UchuPalau_1, ISU_UchuPalau_2; contact: Geoffrey Clark)

Excavations of ~4 m2 in Uchularois Cave by Masse (1989) (*31*)recovered an assemblage of pig bones totalling 110 identified specimens. Radiocarbon dates from Uchularois Cave have been recently revised and suggest that midden deposition began at AD 900, with site abandonment by AD 1450 (*32*). ISU_UchuPalau_1 was radiocarbon dated to ~1,300calAD (OxA-44064).

Pamayan, Philippines (1420–1630 AD; ISU_Philippines_3; contact: Peter Bellwood)

Pamayan, Savidug village, Savidug Island, Batanes Islands, northern Philippines (Late prehistoric, overlapping with early Spanish period). The Pamayan shell midden lies at the back of Savidug village, exposed by a road cutting along the base of a limestone hill. A surface presence of red-slipped pottery indicated the existence of a potentially interesting site, so two small excavation sections of 60 by 80 cm were excavated 16 m apart back into the section exposed in the road cutting, exposing shell midden to a maximum depth of 105 cm beneath the topsoil (*33*). However, the recency of the Pamayan midden was rendered obvious by the occurrence of a few imported Chinese ceramic sherds down to 95 cm depth. A C14 date of 418±41 uncal. BP (WK 13091) from the base of the cultural layer indicates that the midden overlaps in its age of formation with the arrival of Spanish missionaries on the island, although there is a strong possibility that the archaeological materials have migrated downslope from a former site on top of the hill, a circumstance which makes the precise date of the pig bone sample uncertain. The pig bone was therefore dated using radiocarbon dating to 1475 calAD (OxA-43990).

Savidug Dune Site, Philippines (600-1000 BP; ISU_Philippines_1; contact: Peter Bellwood)

Savidug Dune Site, Savidug village, Savidug Island, Batanes Islands, northern Philippines. The Savidug Dune Site has two stratigraphic and cultural phases of usage (*33*). An upper cultural layer dated c.1000 to 600 BP, with imported ceramics and iron, hence overlapping in date with the formation of the Pamayan shell midden. A lower cultural layer dated c.3200 to 2000/1500 BP, which can be divided into an earlier and a later phase. The earlier phase is associated with rare examples of circle-stamped pottery, dated 3200 to 2700 BP in this site and from parallels at Sunget on Batan Island and Anaro on Itbayat Island. The later phase, dated c.2700 to 2000/1500 BP, is associated with jar burial and Taiwan nephrite.

Radiocarbon dating

Two samples were radiocarbon dated at the Oxford Radiocarbon Accelerator Unit (ORAU) The dates were calibrated on OxCal using the IntCal20 curve (*34*):

| Sample ID | Site, country | Uncalibrated radiocarbon age BP | Error (+/-) | Calibrated age BP | Calibration curve | Lab code |
|---|---|---|---|---|---|---|
| ISU_UchuPalau_1 | Uchul a Rois, Palau | 1370 | 18 | 1308-1279 | IntCal20 | OxA-44064 |
| ISU_Philippines_3 | Pamayan, Philippines | 424 | 16 | 511-475 | IntCal20 | OxA-43990 |

Museum specimens

Museum samples for DNA were collected from the AMNH (American Museum of Natural History, New York, USA), the Australian Museum (Sydney, Australia), FMNH (Field Museum of Natural History, Chicago, USA), the Long An Museum (Vietnam), the NHM (Natural History Museum, London, UK), the NMS (National Museum of Scotland, Edinburgh, Scotland), the Naturalis (Naturalis Leiden, The Netherlands), the OUM (Oxford University Museum, Oxford, UK), and the USNM (Smithsonian National Museum of Natural History, Washington DC, USA). More information for these samples can be obtained from Supplementary Data S1.

Archeological and Museum specimen extraction and library

DNA extraction

Samples were extracted in two different ancient DNA facilities - Durham University and the University of Oxford - using conventional set-up for ancient and degraded genomic material (strictly isolated pre- and post-PCR rooms, use of full PPE with overshoes and double set of gloves [nitrile and latex], systematic extraction and library blanks was added to each batch of 12 samples. All equipment and work surfaces were cleaned before and after each use with a dilute solution of bleach (5-10%) followed by water and ethanol (99%). Pipettes and plastic racks were UV-irradiated in a crosslinker (254 nm wavelength) prior to and after use and/or decontaminated using “DNA AWAY” (Fisher Scientific SAS) or bleach followed by purified water. In both laboratories the ancient pig remains were prepared for DNA extraction by removing approximately two a millimetres layer of the outer bone surface by abrasion using a Dremel drill with clean cut-off wheels (Dremel no 409), targeting compact cortical bone or dentine. The bone was then pulverised in a Micro-dismembrator (Sartorious-Stedim Biotech), followed by collection in 15 mL Grainer tubes (Durham) or in 2 mL ependorf tubes (Oxford). Milling containers and grinding balls were subsequently suspended and cleaned in 1% virkon (Durham) or “DNA AWAY” (Oxford) and rinsed in absolute ethanol.

In Durham, the samples were extracted using the protocol described in (35). Bone powder (100-400 mg) was digested in 0.425 M EDTA, 0.05% SDS, 0.05 M Tris-HCI and 0.333 mg/mL proteinase K and incubated overnight (18-24 hours) on a rotator at 50 °C, or until fully dissolved. The digestion buffer, excluding proteinase K, was UV-irradiated (254 nm wavelength) for an hour in a dedicated cross-linker prior to use. 2mL of extract solution was then concentrated in a Millipore Amicon Ultra-4 30 KDa MWCO (Millipore) to a final volume of 100µL. The concentrated extract was purified using silica spin-columns (QIAquick PCR Purification Kit, Qiagen) following manufacturers recommendations, except that the final elution step was performed twice to produce a final volume of 100µL. One in five or one in ten negative extraction controls were performed alongside the ancient bone samples.

In Oxford, we used 80-120 mg of bone or dentine powder that was first pre-digested in 1 mL a buffer made of EDTA and proteinase K for 1h on a rotator at 37 °C, to remove extracellular DNA and some contaminants ((36). After centrifugation, this first lysis was removed and stored in the freezer. The pellets were then fully digested in 1.8 mL of the same digestion buffer, incubated overnight on a rotator at 50 °C. The DNA was extracted from the resulting lysis using a modified (37) protocol. In brief, the 1.8 mL of solution were mixed to 50 ml of the extraction buffer 0.5 M EDTA, pH 8.0 to a final concentration of 0.45 M, 10 mg/ml Proteinase K to a final concentration of 0.25 mg/ml and Tween 20 to a final concentration of 0.05%) and centrifuged through a silica spin column (Qiagen) using an extender. Once bound to the silica filter, the DNA was purified using the commercially provided buffer (PE) and eluted in 42 μL of TET.

Library building and sequencing

The libraries were built in Oxford from the DNA extracts obtained in Durham or Oxford, without distinctions, using the Blunt End Single Tube (BEST) protocol (38). The amplification of the indexed DNA was optimised using quantitative PCR to estimate the best number of cycles. As expected, the number of required cycles varied largely between archaeological and museum samples. In some instances, we performed multiple parallel amplifications to increase the resulting amount of endogenous DNA. Sequencing of selected libraries was performed on a NovaSeq S4 platform (2×150 bp) at Novogene and Macrogen.

Whole genome enrichment and sequencing

Libraries from the following individuals: ASU_AnSon_1, ASU_ManBac_1, ASU_Taiwan_1, ISU_Philippines_1, ISU_Philippines_3, ISU_UchuPalau_1, ISW_Moluccas_1, POU_Vanuatu_3, POU_Vanuatu_5 were subjected to whole genome enrichment (WGE). Genomic DNA from 3 individuals: i.e. a wild boar from Japan (Sus scrofa), a European wild boar (Sus scrofa), and a bearded pig (Sus barbatus) from the Department of Animal Sciences, University of Illinois, was provided to Daicel Arbor Bioscience for baits construction. The gDNA was visually inspected for color and viscosity and the total DNA was quantified via a spectrofluorimetric assay. WGE baits were prepared from equal masses of the 3 gDNA samples.

Libraries were provided to Daicel Arbor Bioscience for capture. They were quantified via quantitative PCR (qPCR). Based on the quantification results (ranging from 1.9ng-445.6ng), the libraries were reamplified for 4-8 cycles to increase the mass, targeting approximately 1000ng per library. Amplified libraries were quantified again via qPCR. For libraries prepared from the same individual, equal masses of each library were pooled for capture.

Each capture pool was dried down to 7 uL by vacuum centrifugation. Captures were performed following the myBaits v5.02 protocol using the custom Pig WGE baits with an overnight hybridization and washes at 60C. Post-capture, the reactions were amplified for 10-18 cycles and were quantified with a spectrofluorimetric assay. A second round of capture was performed in the same manner. The post-capture material was visualized using the TapeStation 4200 (Agilent) platform with a High Sensitivity D1000 tape. Captures that contained adapter artifact underwent gel excision to remove the artifact. The captures were pooled in approximately equimolar ratios. Samples were screen sequenced on the Illumina NovaSeq 6000 platform on a S4 PE150 lane.

Modern samples

Papua New Guinea, Bhutan, and Sri Lankan (contact: Jaime Gongora)

DNA samples from Bhutanese (SAD_Bhutan_1-4; SAW_Bhutan_6-11), Nepalese (ASD_Nepal_1-3), and PNG (PNU_PNG_6-14) pigs were extracted from whole blood using the

QIAamp® DNA Blood Mini Kit; and from hair samples using the Blood and Body Fluid Spin Protocol. DNA from Sri Lankan pigs (SAD_SirLanka_1-4; SAW_SirLanka_5) was extracted using the salting method as described by Miller et al. (39). DNA concentration was determined using ethidium bromide staining and a NanoPhotometer, and ranged from 25-200 ng/µl. All sampling and DNA extraction were done in the respective countries in collaboration with relevant institutions.

With the exception of seven samples (described below), libraries were prepared by the Australian Genome Research Facility (AGRF) as a commercial service using the Illumina Nextera DNA Flex library preparation protocol. Library quality was assessed using a TapeStation and qPCR, and sequenced by AGRF on a NovaSeq 6000 S4-300 flow cell, v1.5 chemistry.

The remaining seven samples (SAW_Bhutan_6-11; SAW_SriLanka_5) were wild boars from Bhutan and Sri Lanka, collected opportunistically from dead, slaughtered, or captured individuals. DNA from these seven samples was potentially degraded because the tissue was not freshly collected. Consequently, aliquots of purified DNA extracted from these samples were sent from the University of Sydney to the Australian Centre for Ancient DNA (ACAD), University of Adelaide, where libraries were created in a clean room using a protocol optimised for fragmented, low-quality DNA. Specifically, DNA was blunt-ended and had truncated Illumina TruSeq adapters ligated following the protocol of (40). Full length Illumina TruSeq adapters (with a unique non-overlapping 7nt i5/i7 index combination per sample) were added via PCR using primer sequences from (41).

Each library was amplified in eight separate 25 µL reactions, each comprising 3 µL of undiluted library, 1 x Platinum Taq DNA Polymerase High Fidelity buffer (ThermoFisher Scientific), 2 mM MgSO4 (ThermoFisher Scientific), 0.25 mM of each dNTP (ThermoFisher Scientific), 0.4 µM of each primer, and 0.2 U of Platinum Taq DNA Polymerase High Fidelity (ThermoFisher Scientific), in laboratory grade water. Cycling conditions for the PCR were as follows: 94 °C for 2 min; 20 cycles of 94 °C for 30 s, 60 °C for 30 s, 68 °C for 40 s; and 68 °C for 10 min. PCR products from each library were pooled and purified using 1.1 x volume AxyPrep (Axygen), washed twice with 80% ethanol, and then resuspended in 30 µL of buffer comprising 10 mM Tris, 0.1 mM EDTA, and 0.05% Tween-20. Library quality was assessed using a Fragment Analyzer (Agilent) and sequenced by AGRF on a NovaSeq 6000 S1-300 flow cell, v1.5 chemistry.

<u>Lanyu pigs</u> (Taiwan; contact: Larry Schook)

DNA was extracted using standard phenol-chloroform techniques. Library preparation and sequencing was carried out at Macrogen (Seoul, South Korea) using an Illumina PCR-free TruSeq library kit (Illumina, Inc) followed by 2x150 bp Paired End analysis on an Illumina NovaSeq instrument.

New Caledonia (contact: Patrick Barriere) and Philippines (contact: Michael Herrera)

DNA extraction was carried out at the PalaeoBARN, University of Oxford. Three hairs per sample were cut into approximately 1 cm long sections and washed in a 0.5% sodium hypochlorite solution. Bleach was removed by rinsing the hairs twice in HPLC grade water followed by two washes in UltraPure DNase/RNase-Free Distilled Water. DNA extraction was carried out using the DNeasy Blood & Tissue Kit (Qiagen), including digestion in Buffer ATL/Proteinase K/DTT solution overnight. Single-stranded DNA sequencing libraries were prepared at the Laboratory of Functional Genome Analysis, Genzentrum, University of Munich

(LMU) Swift BioSciences 1S kit. Sequencing: screening on NextSeq at Genzentrum followed by deep sequencing on a 150 PE NovaSeq 6000 S4 flowcell at Macrogen (Seoul, South Korea).

### Data processing

#### Modern genome

Raw reads from modern genomes from both publicly available samples and newly sequenced, were first processed with fastp (42). Reads were then aligned using BWA-MEM (43), to the Sscrofa11.1 reference genome, duplicates were removed using Picard MarkDuplicates.

#### Historical and ancient genomes

Historical and ancient genomes were processed separately using the nextflow eager pipeline (44) with default parameters except for the following options: --mapper bwamem, and --clip_readlength 20.

### Mitochondrial analyses

#### Mitochondrial genome analysis

We used htsbox (https://github.com/lh3/htsbox) to create a mitochondrial genome majority consensus fasta file based on BAM files using a only reads with a mapping and base quality of 30, and at least 20 base pair long, the first and last 5 base pair were also removed for ancient and historical samples. Fasta files were merged into a single file which was filtered using a custom perl script. Sequences with over 70% missing data were excluded from this analysis, and positions in the alignment with 30% missing data were also removed. This resulted in an alignment of 585 individuals and 15,523bp. A phylogenetic tree was then constructed with RAxML (45) using 100 (fast) bootstrap replicates and a GTRGAMMA substitution model (Figure S1).

#### Published mtDNA control region dataset

Published mitochondrial control region sequences (N =707, see Supplementary Data S2) were aligned using global iterative refinement (G-INS-i) implemented in MAFFT 7.526 (2024/Apr) (46). The alignment was visualised and a 621-bp segment of the control region was selected using Geneious Prime® 2024.0.7 (GraphPad LLC). IQ-TREE multicore version 1.6.12 (47) was used to construct a maximum likelihood tree under a HKY+F+R4 model of evolution, selected using ModelFit (48). Branch support was assessed using 1000 Ultrafast bootstrap psuedo-replicates (49). The bootstrap consensus tree was drawn in FigTree 1.4.5 (https://github.com/rambaut/figtree) and used to assign sequences to one of four haplogroups: Lanyu, Pacific Clade, Europe (European *S. scrofa* in Figure S1), East Asia (Asian *S. scrofa* in Figure S1) , and ISEA (ISEA-endemic *Sus*, i.e. *S. barbatus, S. philipensis/S. cebifrons, S. verrrucosus,* or *S. celebensis* in Figure S1).

### Nuclear analyses

#### Reference panel genotype call

Genotyping was performed using graphtyper v2.7.2 using genomes with at least 5x (500 individuals). The resulting VCF was filtered using the following options:

```
vcffilter -f "ABHet < 0 | ABHet > 0.33 & ABHom < 0 | ABHom > 0.97 & MQ > 30"
bcftools view -f "PASS" -m 2 -M 2 -v snps
```

```
bcftools-1.11/bcftools filter --SnpGap 10 -i "FORMAT/GQ>30" -i "INFO/AC>4"
```

This resulted in a total of ~95M SNPs, including ~25M transversions.

Phasing and imputation

We used SHAPEIT v5 for phasing (27) the reference panel which contains 95M SNPs in 500 individuals. Phasing was performed in 1Mbp sliding windows (100Kb overlap) using the phase_common tool and a recombination map obtained from (50). The resulting BCF were ligated using the ligate tool.

We then used GLIMPSE 2 for imputation (28). We first used the GLIMPSE2_chunk tool to split the phased phase into imputation regions based on the recombination map obtained from (50). Each chunk was then converted into a binary reference format using the GLIMPSE2_split_reference tool. Each BAM file for samples between 1-5x was then imputed separately using GLIMPSE2_phase for each chunk, and chunks were ligated using GLIMPSE2_ligate. The resulting BCFs were then merged with the panel using bcftools.

Imputation accuracy was estimated by first downsampling the BAM files of six individuals to 0.1x, 0.5x, 1x, 2x and 4x using samtools. We then imputed each bams using a reference panel from which these six individuals were removed. We then calculated concordance between "true" genotypes and imputed genotypes (r2) at different minor allele frequency (MAF) using the GLIMPSE_concordance tool (Figure S2). Imputation accuracy was >95% in most individuals from at 2x and MAF>1%. Island Southeast Asian feral individuals from Bali (ISW_Bali_1, ISW_Buru_1), however, only achieved an accuracy of >90% at MAF>5%.

The primary objective of imputation in this analysis is to conduct ancestry deconvolution through local ancestry inference. In addition to testing accuracy of imputation on genotypes, we investigated how imputation influenced our ability to conduct local ancestry inference (LAI) using GNOMIX, to pinpoint genomic segments associated with specific ancestries (see below for details on GNOMIX analysis). We utilized the imputed VCFs derived from downsampled BAM files in the preceding step to conduct a LAI using GNOMIX. For each ancestry, we then compared the regions identified by gnomix in both the full coverage and downsampled data. The results revealed a high degree of overlap between these regions, ranging from 94% to 99% (Figure S3). This suggests that imputation minimally impacted the power of local ancestry inference in this context.

Pseudohaploid call

The 25M transversion SNPs discovered in the 5x data set were then genotyped in historical and ancient genomes not included in the panel using ANGSD (v0.933) with the following options: -dohaplocall 1 -minMinor -doCounts 1 -minMapQ 20 -minQ 20 -minInd 1 -setMinDepth 1 -trim 5. Positions at which the genotype did not match one of the two alleles in the 25M SNP panel were set as missing data. Genotype data were merged using plink v1.9 (51). Samples from the panel were pseudohaplosised using a custom script in the resulting plink file.

Principal component analysis (PCA)

Principal component analysis (PCA) was carried out using emu (52) on pseudohaploid transversions. We filtered SNPs with MAF<5% and kept all samples with >10,000 SNPs covered (576 individuals).

ADMIXTURE

We first ran an unsupervised ADMIXTURE clustering based on 120 pseudohaploid (transversion only) genomes from wild individuals form East Asia, Europe and ISEA to identify individual carrying >99% ancestry from three different clusters: Asian *S. scrofa*, European *S. scrofa* and ISEA-endemic *Sus* (Figure S4). Individuals with at least 99% ancestry from one cluster were then used as a source in a supervised ADMIXTURE analysis (K=3 again), based on 576 individuals.

D and F statistics

Our ADMIXTURE analysis did not possess enough power to split the ancestry of ISEA-endemic Sus individual i.e. *S. celebensis, S. cebifrons, S. verrucosus,* or *S. barbatus*. Therefore, for each ISEA individual that possessed both Asian S. scrofa and ISEA-endemic Sus based on our ADMIXTURE analysis we first used D-statistic to assess which ISEA-endemic Sus best matched their non-scrofa ancestry. To do so we first validated that each individual yielded significant D-statistics of the form D(Babyrousa outgroup, ISEA-endemic Sus; Test, Asian S. scrofa), where ISEA-endemic Sus represent any genome of *S. celebensis*, *S. cebifrons*, *S. verrucosus*, or *S. barbatus* and Asian *S. scrofa* any genome of individuals that possess >99% Asian *S. scrofa* ancestry in our ADMIXTURE analysis.

We then ran all combinations of D-statistics of the form D(B. babyrousa, Test; ISEA-endemic-Sus-Y, ISEA-endemic-Sus-Z), where Test is the individual possessing S. scrofa and ISEA-endemic Sus ancestry, and ISEA-endemic-Sus-Y, and ISEA-endemic-Sus-Z represent all combinations of ISEA-endemic Sus individual i.e. *S. celebensis, S. cebifrons, S. verrucosus,* or *S. barbatus*. We choose the individual species with the highest D value as the source of ISEA-endemic *Sus* ancestry ancestry which was used for the plot in Figure 2.

Admixture graphs

We suspected that Western Indonesian mixed *S. scrofa* / ISEA-endemic *Sus* (e.g. from Sumatra and nearby islands) possess S. scrofa ancestry that is poorly represented by our panel. To address this, we constructed admixture graphs using Admixtools2 (https://github.com/uqrmaie1/admixtools), to assess the phylogenetic placement whether the S. scrofa ancestry found in these individuals is nested within Asian *S. scrofa* or a sister group to Asian and European *S. scrofa* (Figure S5).

We constructed “skeleton graphs”, tailored to for each *S. scrofa* ISEA genome, informed by the D-statistic pipeline above. For individuals with no detectable I ISEA endemic-Sus ancestry, we constructed admixture graphs containing a *B. babyrousa* as an outgroup, an individual with Asian S. scrofa ancestry, an individual with European *S. scorfa* ancestry, and an individual with South Asian *S. scrofa* ancestry (i.e. Sri Lanka or India). The graph was created using the find_graphs function (plusminus_generations = 5, stop_gen = 50, stop_gen2 = 10). We consistently identified the same topology: (Babyrousa, (South Asian, (Asian, European)))

For individuals with detectable ISEA endemic-Sus ancestry we carried out the same as above, but included a genome corresponding to the relevant non-scrofa species (i.e. the species that was the best proxy for the ISEA endemic-Sus ancestry based on D-statistics). This also consistently identified the same topology: (Babyrousa, (ISEA endemic-Sus, (South Asian, (Asian, European))))

We then added each test genome (i.e. *S. scrofa* genome from ISEA) to its corresponding skeleton graph from 1a or 1b, above, in two different locations to create two admixture graphs for each test genome:

- basic graph 1a/1b-Asian: On the Asian branch, i.e. (Babyrousa, (South Asian, ((Asian, test), European))), 1a; or (Babyrousa, (ISEA endemic-Sus, (South Asian, ((Asian, test), European)))), 1b
- basic graph 1a/1b-Sister: On the basal branch, i.e. (Babyrousa, (ISEA endemic-Sus, (South Asian, (test, (Asian, European))))) or (Babyrousa, (South Asian, (ISEA endemic-Sus, (Asian, European))))

For the basic graphs for individuals with detectable non-scrofa ancestry (basic graph 1b-Asian & 1b-Sister), we added an admixture edge between ISEA endemic-*Sus* and our test genome.

For each individual (test), we then assessed whether the Asian or Sister graph was the best fit, using 100 bootstrap-resampled graph fits, after computing out of sample scores to fairly compare models of different complexity to each other.

The ratio of scores for the two admixture graphs were plotted in Figure S5, as:
*(Score_Asian - Score_Sister ) / Max[Score_Asian, Score_Sister]*

<u>Local ancestry reconstruction (GNOMIX)</u>

Local ancestry reconstruction was performed using GNOMIX (29). Wild individuals with Asian *S. scrofa*, European *S. scrofa* and non-scrofa ancestry that were used as a source in the supervised ADMIXTURE above were used as reference for this analysis.

We trained two models, each with two sources:

- *S. scrofa* (using European *S. scrofa*) and non-scrofa
- European *S. scrofa* and Asian *S. scrofa*.

For model 1, we did not use Asian S. scrofa to discriminate between scrofa and non-scrofa ancestry as previous studies have suggested that mainland Southeast Asian wild boar (*S. scrofa*) possess some small degree of ISEA-endemic Sus ancestry (18, 19).

We ran GNOMIX with default settings except for the time of admixture used for validation (generation since admixture: 2, 4, 6, 8, 12, 16, 24, 48). Both models achieved training accuracy and validation accuracy > 99%. Local ancestry reconstruction was then performed on *S. scrofa* individuals from Asia that were either phased (>5x) or imputed (>1x) and which were showed to possess either European *S. scrofa* or non-scrofa ancestry (non possessed both), based on the ADMIXTURE analysis, using the appropriate model (model 1 for non-scrofa ancestry, model 2 in case of European *S. scrofa* ancestry). See the section on imputation above for details about the results of the test for accuracy of imputation for LAI.

<u>Ancestry deconvolution</u>

We used the GNOMIX results to identify regions of Asian *S. scrofa* ancestry in the genome of *S. scrofa* individuals with mixed ancestry, i.e. possessing either non-scrofa and Asian *S. scrofa* (GNOMIX model 1) or European S. scrofa and Asian scrofa ancestry (GNOMIX model 2). To do so, we selected regions which were identified with high confidence (probability >99%) by

GNOMIX as either homozygous *S. scrofa* (model 1) or homozygous Asian *S. scrofa* (model 2). For each mixed individual, we extracted these regions from the pseudohaploid PLINK file used for PCA, ADMIXTURE, and D-statistics analyses described above.

We then calculated admixture fraction using f4 ratio to assess the proportion of non Asian *S. scrofa* ancestry left in the genomes of mixed individuals after deconvolution (Figure F4-deconvolution). European ancestry in most (14/16) individuals from New Caledonia, Hawai, and Papua New Guinea decreased from ~50-25% to 0-5%. European ancestry in one individual from New Caledonia, however, remained at ~10%, at ~20% in an individual from Papua New Guinea. European ancestry in Lanyu pigs decreased from ~25% to 0-1%, while European ancestry in domestic pigs from the Philippines decreased from 30-70% to 0-15%.

Non-scrofa ancestry in individuals with lower *S. scrofa* ancestry such as pigs from Sibago or pigs East of the Wallace line, including Flores, Timor, and Halmhaera decreased from between ~80-40% to below 5%. The non-scrofa ancestry in wild boars from Western Indonesia (West of the Wallace line), including Sumatra and nearby islands, however, decreased from between 14-5% to ~5%. The lower success of deconvolution in these individuals can likely be explained by ancient admixture processes (18) which would be found in smaller blocks of non-scrofa ancestry across the genome that are undetectable by GNOMIX. In contrast, due to its recent introduction in the region, European S. scrofa ancestry is found in larger blocks (Figure 3B), which are more easily detectable.

<u>outgroup f3</u>

We calculated outgroup-f3 statistics using calc-f3 from the structf4 package (53) and babirusa as an outgroup. The goal of this analysis is to assess the degree to which Lapita pigs from Melanesia share drift with other populations of S. scrofa. To do so we first chose a 2,500years old ancient Vanuatu genome (POU_Vanuatu_3; ~0.7x) as reference and computed all pairwise outgroup-f3 involving this individual.

We found that feral pigs from Papua New Guinea (e.g. ISW_PNG_18; ~2.2x) shared the most drift with this individual. ADMIXTURE analysis of ISW_PNG_18 did not detect any evidence of European ancestry, making it a good proxy for Lapita pig ancestry. We used this individual as reference in subsequent outgroup-f3 analyses due to its higher coverage than POU_Vanuatu_3 to limit issues arising when comparing multiple individuals with low breadth of coverage, particularly after ancestry deconvolution. We then computed all pairwise outgroup-f3 involving ISW_PNG_18 using both the full and deconvoluted dataset.

<u>Admixture time (DATES)</u>

We used DATES (26) to estimate time of admixture in admixed Island Southeast Asian individuals using the recombination map obtained from (50) and the following parameters:

*binsize: 0.001*
*maxdis: 1*
*jackknife: YES*
*qbin: 10*
*runfit: YES*
*afffit: YES*
*lovalfit: 0.45*

The same individuals used to train the GNOMIX models were used as source for this analysis. We used a generation time of 3 years to convert into years. Results were considered significant at first if Z-score > 2 and normalized root-mean-square deviation (NRMSD) < 0.7. We first assessed the accuracy of DATES using five European domestic individuals with known dates of admixture with Asian *S. scrofa* (as a result of 19th century improvement of pig stock in Europe). We modelled these individuals using the same source as for the ADMIXTURE and GNOMIX analyses.

All results were significant and visual inspection of the exponential distribution fitted and the covariance decay curve suggested a good fit (Figure S6). The recovered admixture proportion and admixture dates admixture dates fitted well with the known history of these breeds (admixture date in the middle of the 19th century (54)):

| **Sample ID** | **Sources** | **EUW** | **ASW** | **Mean time (generation)** | **SE** | **Z** | **NRMSD** | **Mean time (years)** |
|---|---|---|---|---|---|---|---|---|
| EUD_Duroc_1 | EUW-ASW | 0.78 | 0.22 | 55.5 | 15.0 | 3.7 | 0.083 | 1833 |
| EUD_Landrace_11 | EUW-ASW | 0.784 | 0.216 | 39.5 | 8.5 | 4.6 | 0.058 | 1881 |
| EUD_Pietrain_10 | EUW-ASW | 0.772 | 0.228 | 46.8 | 8.1 | 5.8 | 0.069 | 1860 |
| EUD_Tamworth_3 | EUW-ASW | 0.831 | 0.169 | 59.9 | 14.4 | 4.1 | 0.07 | 1820 |
| EUD_Yorkshire_19 | EUW-ASW | 0.758 | 0.242 | 53.7 | 7.8 | 6.9 | 0.073 | 1839 |

We modelled individual from Oceania, Melanesian and the Philippines using the same approach:

| **Sample ID** | **Sources** | **EUW** | **ASW** | **Mean time (generation)** | **SE** | **Z** | **NRMSD** | **Mean time (years)** |
|---|---|---|---|---|---|---|---|---|
| ISD_Lanyu_1 | EUW-ASW | 0.351 | 0.649 | 6.221 | 1.987 | 3.131 | 0.137 | 1981 |
| ISD_Lanyu_2 | EUW-ASW | 0.29 | 0.71 | 8.63 | 7.832 | 1.102 | 0.077 | 1974 |
| ISD_Lanyu_3 | EUW-ASW | 0.319 | 0.681 | 3.147 | 6.363 | 0.495 | 0.094 | 1991 |
| ISD_Lanyu_4 | EUW-ASW | 0.342 | 0.658 | -2.435 | 10.307 | -0.236 | 0.136 | 2007 |
| ISD_Lanyu_5 | EUW-ASW | 0.308 | 0.692 | -5.063 | 13.104 | -0.386 | 0.115 | 2015 |
| ISU_Philippines_10 | EUW-ASW | 0.341 | 0.659 | 18.075 | 2.469 | 7.321 | 0.087 | 1946 |
| ISU_Philippines_11 | EUW-ASW | 0.304 | 0.696 | 38.298 | 15.299 | 2.503 | 0.107 | 1885 |
| ISU_Philippines_12 | EUW-ASW | 0.647 | 0.353 | 16.761 | 11.669 | 1.436 | 0.073 | 1950 |
| ISU_Philippines_13 | EUW-ASW | 0.739 | 0.261 | 24.639 | 3.955 | 6.229 | 0.091 | 1926 |
| ISU_Philippines_14 | EUW-ASW | 0.696 | 0.304 | 36.515 | 11.697 | 3.122 | 0.088 | 1890 |
| ISU_Philippines_15 | EUW-ASW | 0.753 | 0.247 | 18.08 | 9.237 | 1.957 | 0.078 | 1946 |
| ISU_Philippines_4 | EUW-ASW | 0.673 | 0.327 | 6.702 | 5.378 | 1.246 | 0.166 | 1980 |

| | | | | | | | | |
|---|---|---|---|---|---|---|---|---|
| ISU_Philippines_5 | EUW-ASW | 0.54 | 0.46 | 18.577 | 8.305 | 2.237 | 0.077 | 1944 |
| ISU_Philippines_6 | EUW-ASW | 0.739 | 0.261 | 10.952 | 16.668 | 0.657 | 0.109 | 1967 |
| ISU_Philippines_7 | EUW-ASW | 0.648 | 0.352 | 40.351 | 11.847 | 3.406 | 0.087 | 1879 |
| ISU_Philippines_8 | EUW-ASW | 0.475 | 0.525 | 36.789 | 9.636 | 3.818 | 0.067 | 1890 |
| ISU_Philippines_9 | EUW-ASW | 0.791 | 0.209 | 63.41 | 11.33 | 5.597 | 0.05 | 1810 |
| ISW_Marquesas_1 | EUW-ASW | 0.639 | 0.361 | 34.074 | 9.877 | 3.45 | 0.096 | 1898 |
| ISW_Morotai_1 | EUW-ASW | 0.177 | 0.823 | 30.923 | 6.229 | 4.964 | 0.074 | 1810 |
| ISW_Morotai_2 | EUW-ASW | 0.186 | 0.814 | 30.347 | 6.181 | 4.91 | 0.072 | 1812 |
| ISW_PNG_20 | EUW-ASW | 0.208 | 0.792 | 16.317 | 5.572 | 2.929 | 0.091 | 1922 |
| ISW_Sumba_1 | EUW-ASW | 0.453 | 0.547 | 17.65 | 2.946 | 5.991 | 0.189 | 1877 |
| PNU_PNG_10 | EUW-ASW | 0.424 | 0.576 | 26.165 | 4.892 | 5.349 | 0.058 | 1922 |
| PNU_PNG_11 | EUW-ASW | 0.421 | 0.579 | 7.609 | 8.702 | 0.874 | 0.108 | 1977 |
| PNU_PNG_12 | EUW-ASW | 0.544 | 0.456 | 25.227 | 3.32 | 7.599 | 0.117 | 1924 |
| PNU_PNG_13 | EUW-ASW | 0.571 | 0.429 | 25.036 | 8.263 | 3.03 | 0.076 | 1925 |
| PNU_PNG_14 | EUW-ASW | 0.482 | 0.518 | 17.289 | 8.536 | 2.025 | 0.078 | 1948 |
| PNU_PNG_6 | EUW-ASW | 0.487 | 0.513 | 6.311 | 17.201 | 0.367 | 0.101 | 1981 |
| PNU_PNG_7 | EUW-ASW | 0.334 | 0.666 | 21.447 | 10.853 | 1.976 | 0.077 | 1936 |
| PNU_PNG_8 | EUW-ASW | 0.418 | 0.582 | 30.253 | 13.904 | 2.176 | 0.073 | 1909 |
| PNU_PNG_9 | EUW-ASW | 0.488 | 0.512 | 14.085 | 4.373 | 3.221 | 0.13 | 1958 |
| POW_Hawaii_1 | EUW-ASW | 0.585 | 0.415 | 40.041 | 8.144 | 4.916 | 0.078 | 1817 |
| POW_NewCaledonia_1 | EUW-ASW | 0.57 | 0.43 | 23.775 | 4.149 | 5.73 | 0.081 | 1942 |
| POW_NewCaledonia_2 | EUW-ASW | 0.462 | 0.538 | 33.71 | 7.245 | 4.653 | 0.081 | 1912 |
| POW_NewCaledonia_3 | EUW-ASW | 0.544 | 0.456 | 41.317 | 8.477 | 4.874 | 0.071 | 1889 |
| POW_NewCaledonia_4 | EUW-ASW | 0.544 | 0.456 | 30.252 | 9.299 | 3.253 | 0.082 | 1922 |
| POW_NewCaledonia_5 | EUW-ASW | 0.587 | 0.413 | 23.563 | 15.314 | 1.539 | 0.08 | 1942 |
| POW_NewCaledonia_6 | EUW-ASW | 0.45 | 0.55 | 31.695 | 7.888 | 4.018 | 0.045 | 1918 |

Most significant (Z>2) results points to dates in from the early 18th century to the mid 20th century. We attempted to estimate time of admixture between *S. scrofa* and endemic-Sus in Western Indonesia populations but visual inspection of the exponential distribution fitted and the covariance decay curve suggested a poor fit (Figure S7). This is likely due to the fact that there are only the low number of individual genomes available for each ISEA-Endemic Sus species and possibly that the Asian *S. scrofa* population used here is a poor proxy for the ancestry in these populations;

Geometric Morphometric (GMM)

Data acquisition

Information on the specimens analysed with geometric morphometrics can be found in Data S3. This study used geometric morphometrics to analyze the shape and size of third lower molars. Digital images of the teeth were taken from above (occlusal view) using a standardized protocol (55). Twelve anatomical landmarks and 87 sliding semi-landmarks (Figure S8) were then placed on these images using TPSdig software (56). Coordinates were superimposed using a General Procrustes Analysis (function ProcSym of the R package Morpho (57)). During this procedure the position of the SSL were adjusted using the Procrustes distance (bending = FALSE) and the original configuration was used in all iterations (recursive=FALSE). The centroid size and Procrustes residuals (coordinates after superimposition) were used respectively to assess tooth size and variation.

Predictive Discriminant Analysis

In order to identify the ancient teeth corresponding to *S. scrofa* several regional predictive linear discriminant analyses were performed. The identification was based on a resampling approach based on groups of the same size combined with a data reduction procedure selecting the first components (PC) of a Principal Component Analysis (PCA) that maximise the between group differentiation (58). Among the 100 analyses computed, only the discriminant analyses with the highest correct cross-validation percentages (CVP, jackknife) were retained (upper quartile of the distribution). The specimens of *S. scrofa* included in those identifications originated from China, India, Thailand, Vietnam, Malaysia, Burma, Sumatra and Java. We then classified individual samples from Island Southeast Asia using this approach.

In Sulawesi, the 4 ancient specimens could correspond either to *S. scrofa* (N=97) or *S. celebensis* (N=93). The two modern species have a correct cross-validation of 96% (90% confidence interval CI: 95.7%-96.8%) using the 17 first PCs carrying 91.3% of the total variance. For the prediction, 67 discriminant analyses were retained all with a CVP above 96%. The four specimens were identified as Sus scrofa.

In Sarawak (Borneo), 26 archaeological specimens were compared to 67 *S. barbatus* and 97 *S. scrofa*. The CVP of the two species is 90.7% (CI: 88.7-93.2) based on 12 PCS carrying 85.9% of the total variance. 39 discriminant analyses were retained, all with a CVP above 91%. 13 archaeological specimens were identified as *S. barbatus* and 13 to *S. scrofa*.

For the Philippines, the 48 archaeological specimens were identified as either S. scrofa (N=97) or ISEA endemic Sus (N=38, including *S. cebifrons* (N=8), *S. oliveri* (N=3), and *S. philippensis* (N=27)). The two modern taxa have a CVP of 92.7% (CI: 88.1-97.4%) based on 13 PCs (% of total variance). 36 discriminant analyses were retained resulting in the identification of 31 *S. scrofa* and 17 endemic Sus.

In Flores, 6 archaeological specimens were identified as either *S. celebensis* (N=93) or *S. scrofa* (N=97). The cross validation between the two species was 96.3% (CI: 96.2, 96.8%) based on 13 PCs (% of variance). 97 discriminant analyses were retained, with a CVP above 96.2%, that allowed to identify 5 *S. celebensis* and 1 *S. scrofa*.

The 147 archaeological specimens from mainland South-East Asia (China, Vietnam, Thailand) strongly differ from the 31 from the far east of the distribution (Marqueses, Vanuatu). Between those two groups the correct cross validation is 97% (CI: 92.3-100%) based on the first 10 PCs (83% of total variance). The Wallacean and Pacific specimens exhibited a third lower molar proportionally wider and shorter compared to the more rectangular shape of the specimens from mainland Asia (Figure S9). Based on those PCs and the predictive discriminant analysis approach described above (see (58)) the remaining 102 specimens were identified based on 42 discriminant analyses with a CVP above 98%.

The morphometric proximity between ancient and modern groups from the various regions and taxonomy was visualised using Mahalanobis distances obtained from the 15 first PC scores (88.7% of total variance) that maximise the between group differences (Fig. 1C).

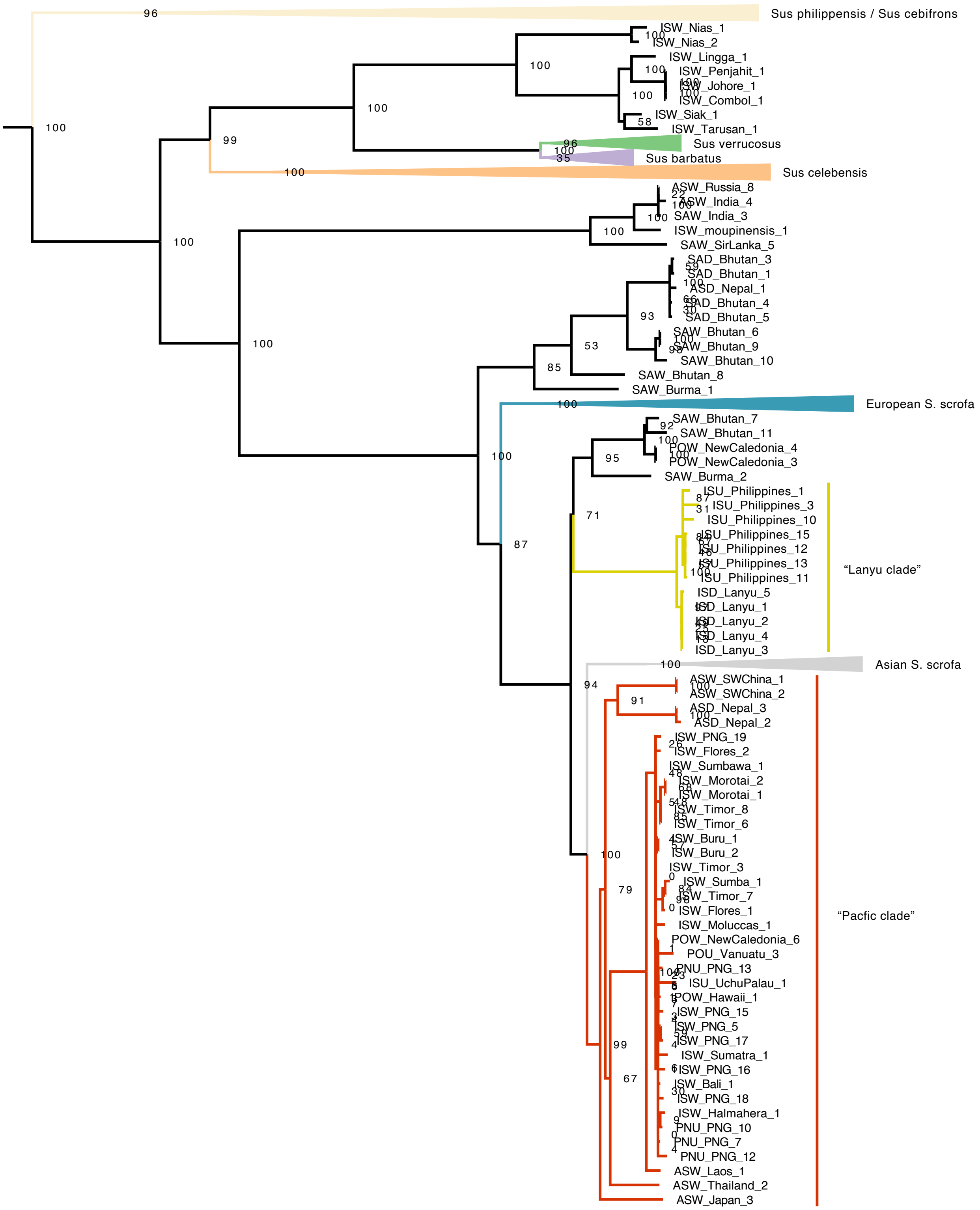


**Fig. S1.**

Maximum likelihood tree based on 585 mitogenomes with bootstrap support.

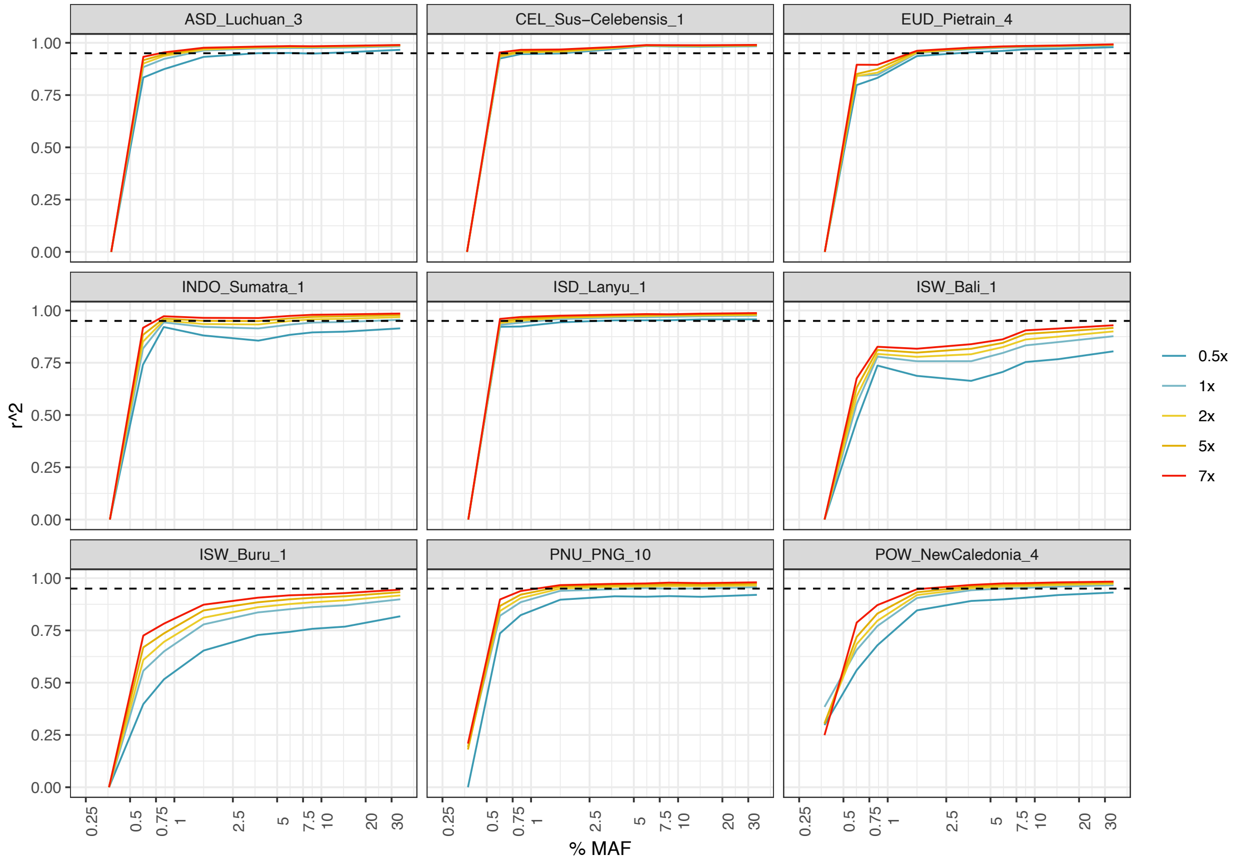


**Fig. S2.**

Imputation accuracy at different coverage. The x-axis represents MAF (minor allele frequency), y-axis represents concordance between "true" genotypes and imputed genotypes (r2) at different downsampled coverage (0.5-7x) as computed by the GLIMPSE_concordance tool.

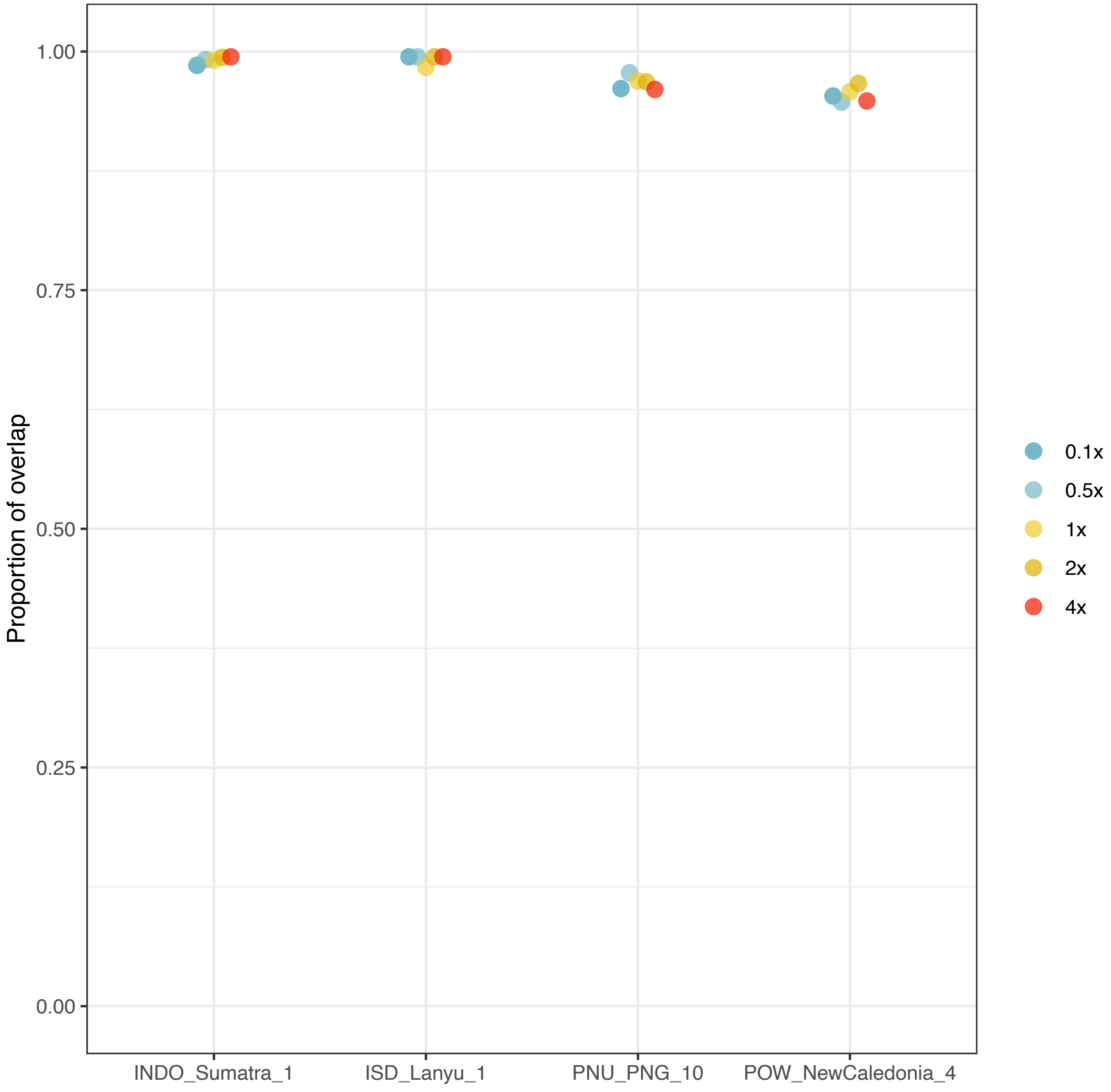


**Fig. S3.**
Accuracy of local ancestry inference across different sequencing coverages. The y-axis shows the proportion of matching ancestry calls by GNOMIX between high-coverage data and downsampled data 0.5x to 4x coverage.

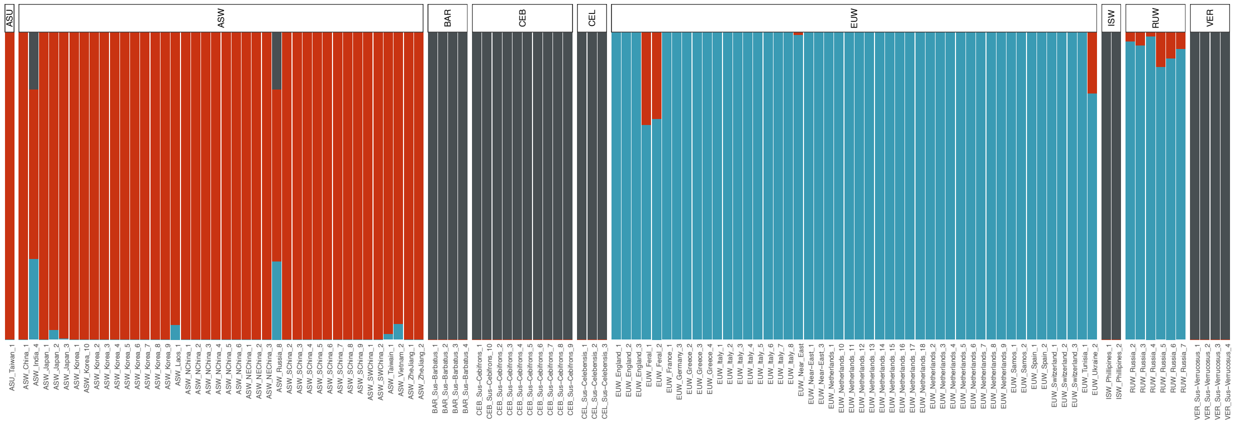


**Fig. S4.**

Results of unsupervised ADMIXTURE analysis using K=3 based 120 wild boar genomes.

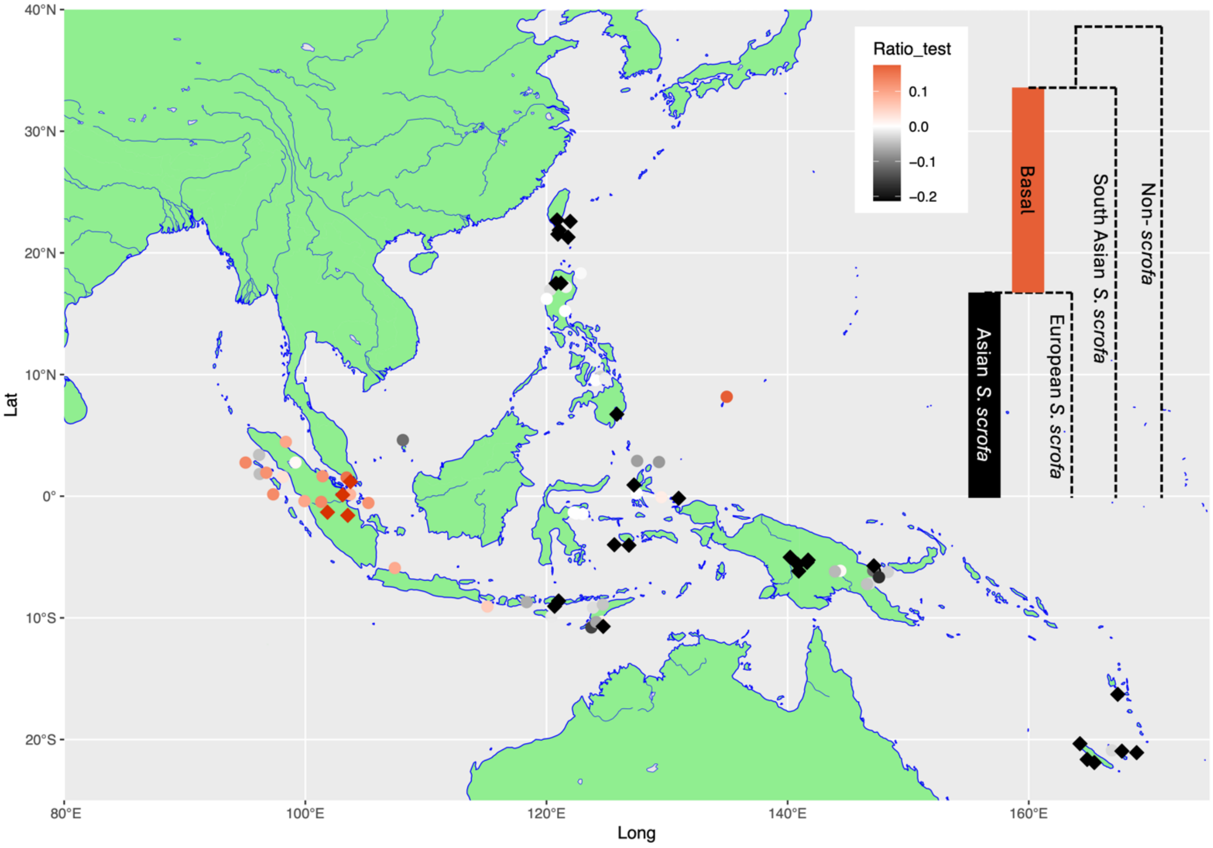


**Fig. S5.**

Figure showing the fit of admixture graphs assuming basal *S. scrofa* ancestry versus Asian *S. scrofa* ancestry for each sample, accounting for non-*S. scrofa* admixture. Models supporting basal *S. scrofa* ancestry are shown in red, and those supporting Asian ancestry in black. Instances when this support is significant at $p<0.05$ is shown by diamonds.

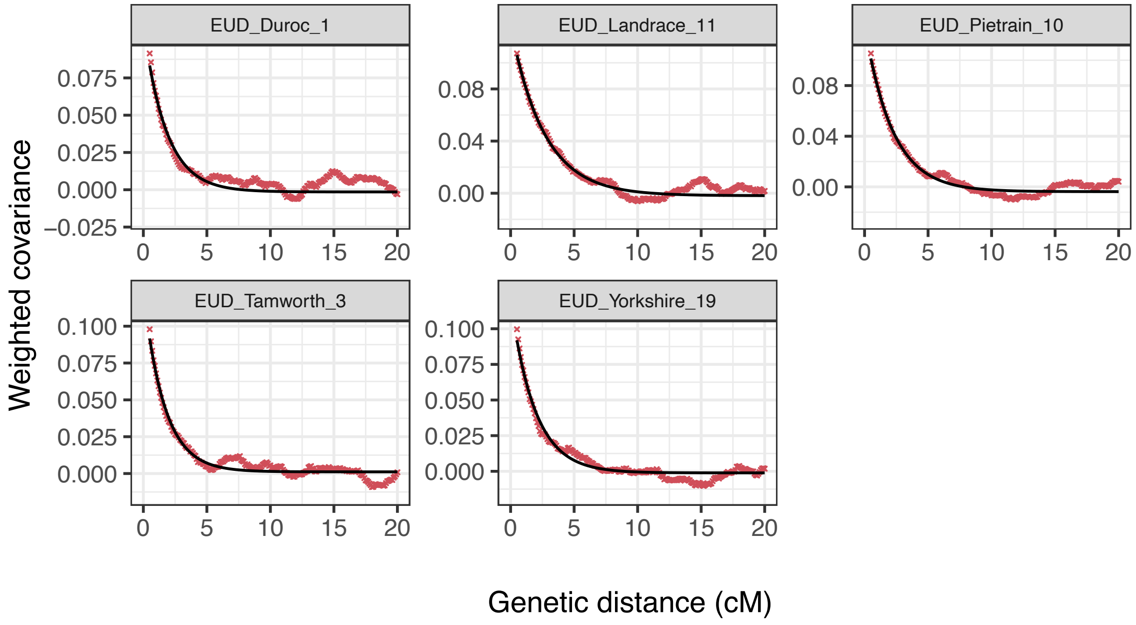


**Fig. S6.**
Example of exponential function fitted by DATES to ancestry covariance in European pigs which are known to possess Asian ancestry.

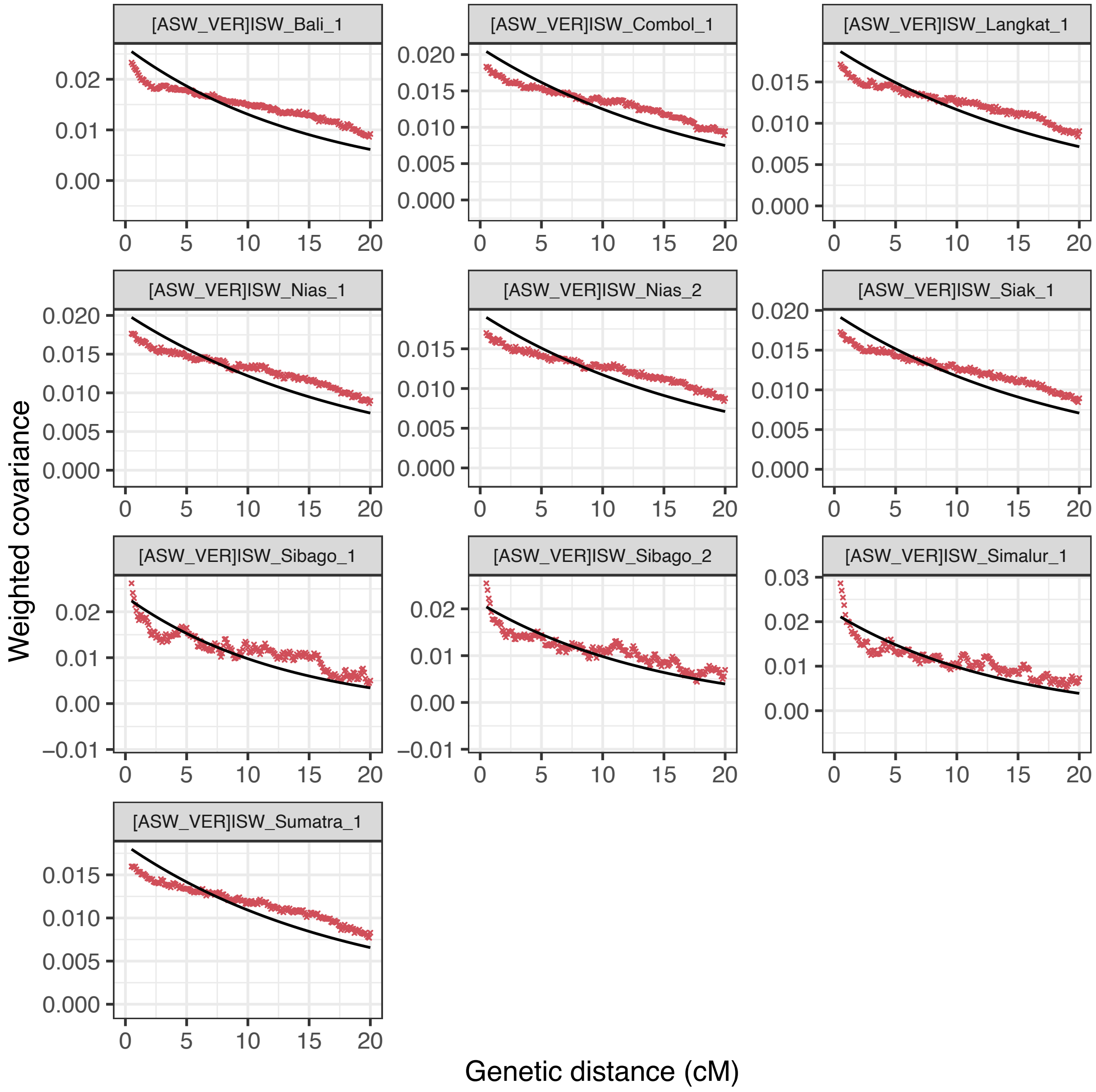


**Fig. S7.**

Exponential function fitted by DATES to ancestry covariance in Asian ancestry which are known to possess *S. verrucosus* ancestry.

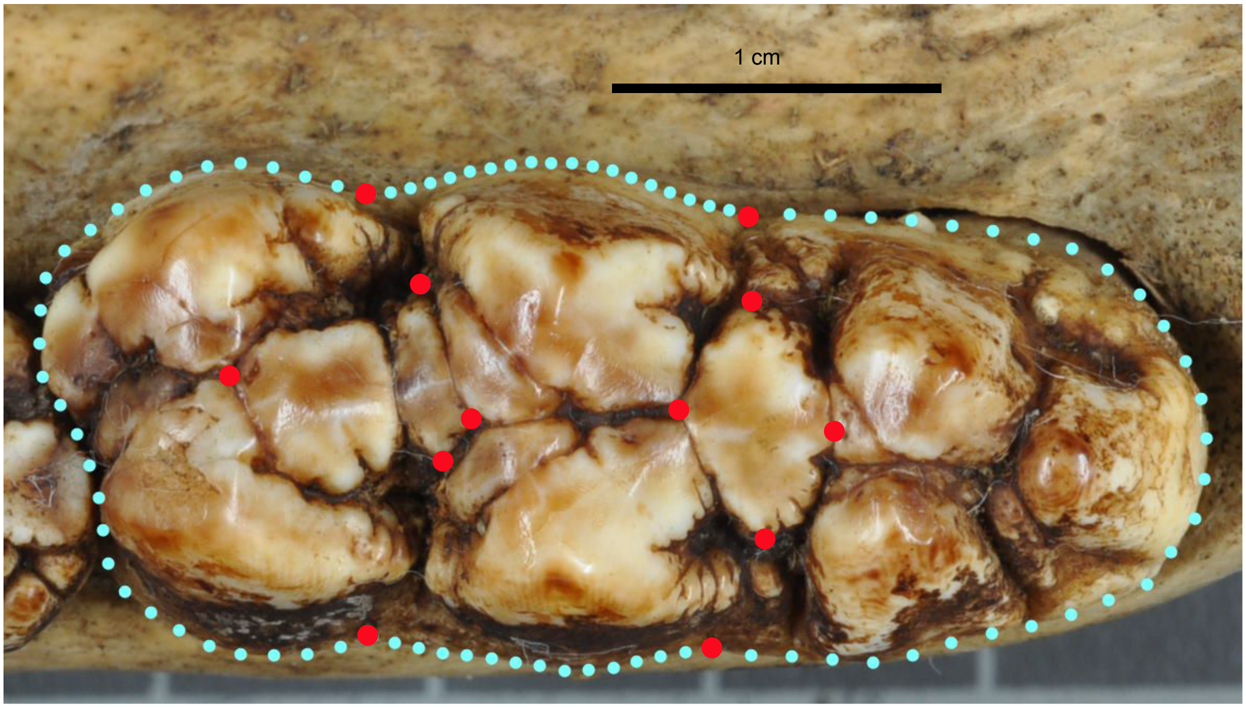


**Fig. S8.**
Position of the 12 2D-landmarks and 87 sliding semi-landmarks on a third lower molar (specimen nhm-69518).

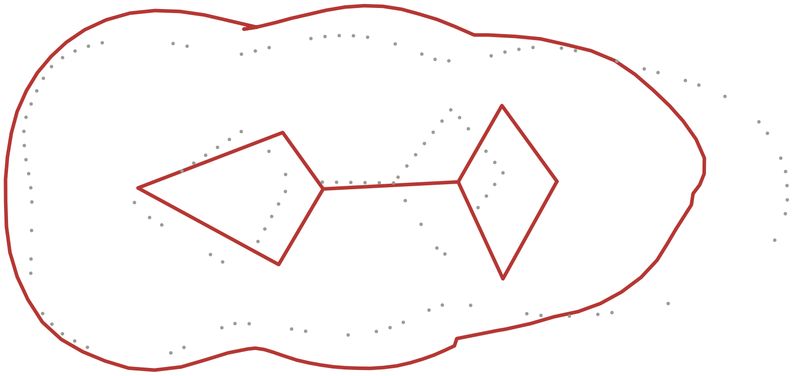


**Fig. S9.**

Molar shape differences between Wallacean and Pacific specimens (N=31) and those from mainland Asia (N=147). Visualisation of the third lower molar shape variation along the discriminant axis between the two groups.

**Data S1. (separate file)**
Information about samples newly sequenced data and publicly available data used in this study including provenance, sampling date, age and sequencing statistics.

**Data S2. (separate file)**
Information about the mitochondrial data used in Figure 2A.

**Data S3. (separate file)**
Information about the GMM specimens.
